# Computational modeling of hormone- and cytokine-dependent proliferation of endometrial cells in 3D co-culture

**DOI:** 10.1101/2025.10.18.682837

**Authors:** Wangui Mbuguiro, Samantha E. Holt, Linda G. Griffith, Juan S. Gnecco, Feilim Mac Gabhann

## Abstract

The endometrium and menstrual disorders, such as endometriosis and adenomyosis, are difficult to study, partly because menstruation depends on interactions between multiple cell types through complex molecular mechanisms. To help understand this system, researchers need humanized experimental and computational models that can interrogate how endometrial cell populations impact each other in both physiological and pathological conditions. Here, we use ordinary differential equations (ODEs) to model changes in the rates of endometrial cell proliferation and death in response to hormones, cytokines, and the specific cell types present. To calibrate this computational model, we used previous-published experimental datasets from a 3D co-culture platform supporting primary human endometrial epithelial organoids and endometrial stromal cells. Our ODE-based model can simulate the size of endometrial epithelial organoids and the density of stromal cells over time under multiple hormone/cytokine conditions in mono- and co-cultures. We further created a second, partial differential equation (PDE)-based model that simulates the diffusion of molecules added to these 3D cultures and their uptake by proliferating endometrial cells using the predicted cell densities from the ODE model as inputs to the PDE simulations. We show that the exposure to hormones and cytokines used in the experiments is reasonably homogenous throughout the 3D culture and identify conditions where this would not be true. Altogether we use these models to quantify the influence of stromal cells on epithelial cell proliferation and *vice versa*, to identify differences across cells from different donors, and to provide a quantitative assessment of experimental designs.

## Introduction

The human endometrium, the inner mucosal lining of the uterus, grows and sheds each month in response to the sex hormones 17*β*-estradiol (“estradiol”, E2, an estrogen) and pregn-4-ene-3,20-dione (“progesterone”, P4) acting on multiple cell types and signaling pathways. Endometrial tissue consists of two layers, the functionalis and the basalis, which are composed of epithelial, stromal, vascular, immune, and stem/progenitor cells (1,2). During menses, the functionalis breaks down and sheds in response to circulating levels of progesterone decreasing alongside estradiol (3). During the proliferative phase of the menstrual cycle, the functionalis regenerates from the basalis as systemic levels of estradiol increase (4). Subsequently, progesterone levels rise in the secretory phase of the cycle, leading to morphologic and functional changes to stromal cells within the functionalis, termed decidualization, to prepare for potential embryonic implantation. The absence of implantation within the endometrium leads to changes in the ovary, triggering a sharp decline in circulating progesterone and estradiol. Menses ensues as the cycle continues (3) (Fig. 1A).

**Figure 1.**
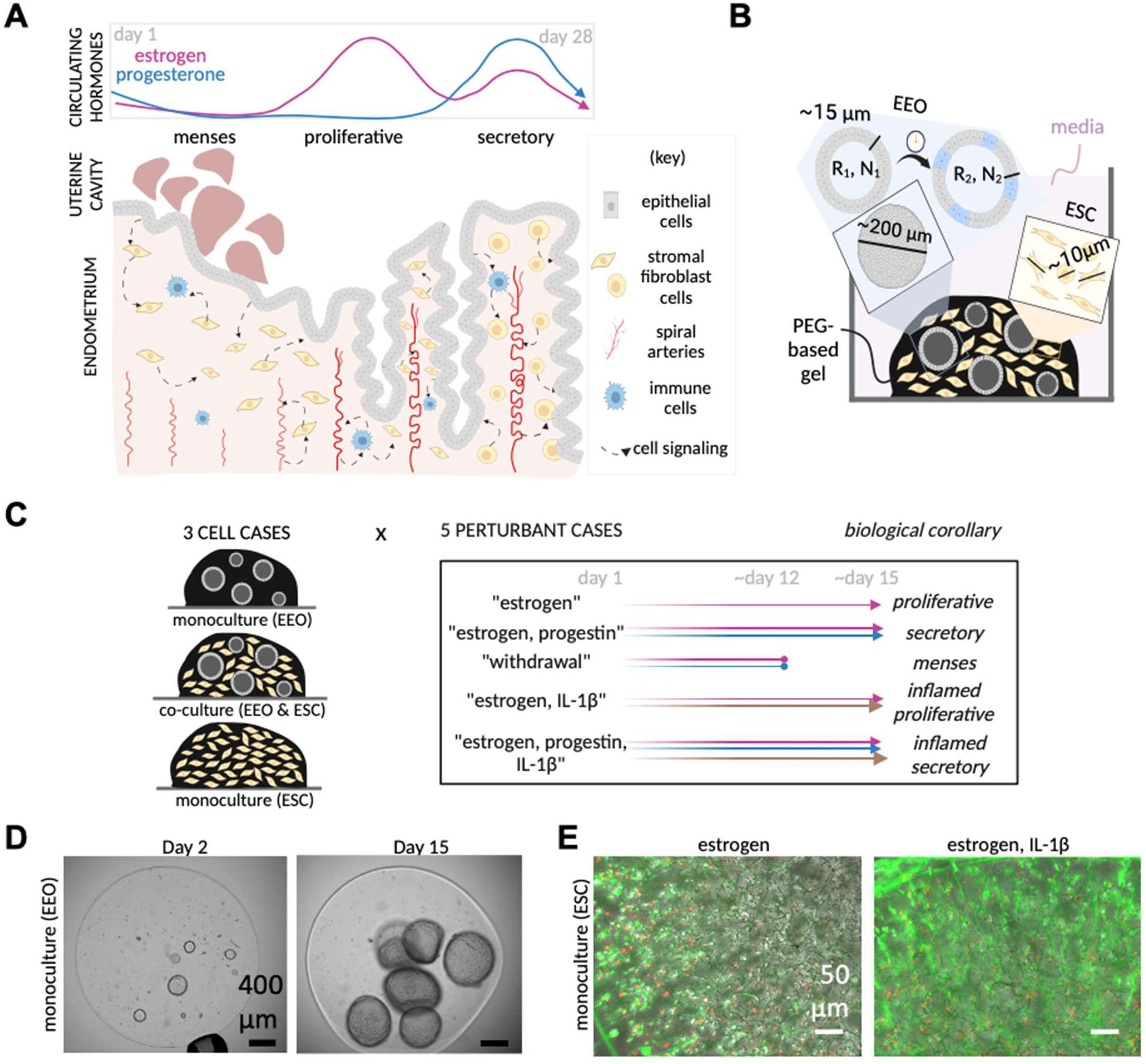
Building and calibrating a computational model of cell proliferation in 3D endometrial co-cultures. **(A)** Overview of Endometrial Tissue. The endometrium undergoes cyclic growth and shedding in response to circulating hormones and cell signaling. **(B)** Two Cell Types in Experimental Co-culture Model. Endometrial epithelial organoids (EEO) are hollow spheres that resemble glands and grow by epithelial proliferation, changing their outer organoid radius (R1, R2) and cell density (N1, N2) over time. Endometrial stromal cells (ESCs) also proliferate but are more dispersed throughout the cell culture. Our computational model predicts how epithelial and stromal cell densities change in response to experimental conditions. **(C)** Experimental Conditions. We use data from fifteen in vitro cell culture conditions. The menstrual phases and inflammation conditions are recapitulated by perturbing in vitro cultures with select hormones and cytokines. **(D-E)** Experimental Data. To calibrate our computational model we used data from a previous study that included measurements of EEO size over time **(D)** and ESC Live/Dead staining at the end of each experiment **(E)**, as described in *Methods*. Figure created in BioRender.

This cyclic process of shedding, regeneration, and decidualization of the functionalis only exists in a small number of non-human species and is not found in the most common animal models used for preclinical research, such as house mice (*Mus musculus*) (5). Furthermore, endometrial tissue dysfunction has been associated with debilitating pain and infertility in patients with disorders such as endometriosis (the growth of endometrial-like cells outside of the uterus) and adenomyosis (the growth of endometrial-like cells within the muscular myometrial layer of the uterus) (6–9). Patients with fibroids (noncancerous smooth muscle tumors within the myometrium) can also experience endometrial abnormalities – including heavy/irregular periods, dysregulated immune signaling, and differences in the mechanical forces from the myometrium that act on the endometrium – which are thought to contribute to the fertility challenges these patients face (10). Because human endometrial tissue function is cyclical, dependent on the interplay of several cell types, and not easily replicated in animal models, there is a need for more humanized *in vitro* and *in silico* models. To further understand healthy endometrial tissues and differences in disease, we need to employ computational and systems biology approaches to develop and analyze results from these humanized *in vitro* models.

To study endometrial tissues *in vitro*, researchers developed 3D culture models using human endometrial cells (as reviewed in (11–13)). By encapsulating cells within a hydrogel matrix which resembles native extracellular matrix, scaffold-containing 3D cultures allow for more realistic cell-cell and cell-matrix interactions which can in turn influence cell genotype and phenotype (14). Over the last 30 years, 3D endometrial cultures have advanced from stromal monocultures (15), to cultures of stroma and epithelia in adjacent compartments (16–19), and now to inter-mixed co-cultures of epithelial and stromal cells (20–22). When cultured in hydrogels with media enriched with specific biological cues, epithelial cells can form spherical organoids, closely resembling the structure and functioning of epithelial glands within the endometrium and endometriosis lesions (23–28). Accordingly, tremendous effort has been made to use 3D cultures to capture patterns in gene expression data to characterize healthy vs. diseased endometrial tissues (26,28) and proliferative vs. secretory endometrial tissues (26,29). These studies have also helped transition from using immortalized cell lines to primary cells, which are thought to be more like *in vivo* endometrial cells (13). Primary cells taken from different donors can also reflect variability in the characteristics of endometrial tissues. Donor-to-donor variability may arise from differences in patient pathologies and co-morbidities (26), the phase of the menstrual cycle at tissue collection (26,30), or other donor characteristics.

Mechanism-based computational modeling can be used to interrogate the influence of individual variability on the dynamics of endometrial tissue growth. The use of mechanism-based modeling to study menstruation has so far been limited to studies of hormone cycling and the pharmaceutical development of hormone-modulating therapies (31–34). To our knowledge, only one mechanistic computational model, developed by Arbeláez-Gomez *et al.* (35), simulates changes to endometrial tissues across the menstrual cycle. That model predicted changes to endometrial tissue volume and shedding in response to cycling levels of estrogen and progesterone using a system of ordinary differential equations (ODEs) (35). Its predictions for changes in endometrial tissue volume closely resembled measurements from an ultrasound study of normally cycling women (36). By including a previous mechanistic computational model of hormone cycling (37) as input to their own model, Arbeláez-Gomez *et al.* (35) were able to predict differences in endometrial volume associated with the use of hormone therapies and validated this with data from a clinical study (38). The computational model showed that differences in estrogen-driven early tissue repair and regeneration may explain incidences of bleeding outside of menses (35). However, this model focused on predicting the volume of endometrium and its menstrual phase; it did not differentiate between individual cell types within the endometrium and has not yet been used to explore menstruation for people with uterine pathologies.

To investigate the influence of different cell types on hormone- and cytokine-dependent endometrial tissue growth, we have created a mechanistic ODE-based model of proliferating endometrial epithelial cells and endometrial stromal cells within 3D co-culture (Fig. 1). To calibrate our model, we simulated experiments from a recent study that cultured donor-matched primary endometrial epithelial organoids (EEOs) and endometrial stromal fibroblast cells (ESCs) in a novel 3D culture system (22). In that study (22), EEOs and ESCs were isolated (≥95% purity), passaged up to 5 times, and cultured in a synthetic hydrogel matrix and exposed to hormones characteristic of the three menstrual cycle phases (proliferative, secretory, and menses). Since the cytokine interleukin-1*β* (IL-1*β*) has been implicated in endometrial pathology (39), its effect on the 3D cultures was also investigated in each menstrual cycle phase (22). We used the experimental datasets to estimate values for the computational model parameters, and we used these parameter values to compare and characterize differences between proliferating endometrial epithelial and stromal cell types alone or in co-culture, as well as differences between tissues from different donors. Knowing that endometrial cells also uptake the hormones and cytokines being added during experiments, we then considered: does the proliferation of endometrial cells in 3D cultures lead to spatial gradients in the molecules added over the course of multi-week experiments? Heterogeneity in the amount of hormone or cytokine that cells are exposed to could lead to differences in cell proliferation, death, and signaling dynamics for cells at different locations in the 3D culture – which should be considered as experimental results are interpreted. To answer this question, we created a partial differential equation (PDE)-based model to predict IL-1*β* distribution across the hydrogel, using the predicted cell densities from our ODE model as inputs to our PDE model. We focused on predicting the distribution of IL-1*β* since it is the molecule of largest size (molecular weight and hydrodynamic radius) added to experiments, so it is most likely to diffuse the slowest. Therefore, IL-1*β* is the most susceptible to forming gradients throughout cultures as it reacts with receptors on proliferating cells. Thus, we modeled IL-1*β* diffusion and reaction in endometrial 3D cultures to predict the heterogeneity in cell exposure to IL-1*β* and further evaluate experimental platform designs.

## Methods

### ODE Model of Endometrial Cell Proliferation and Death

We modeled the density of endometrial epithelial organoid cells (EEOCs) and ESCs in cell culture as continuous functions, dependent on proliferation and death processes, using ordinary differential equations (ODEs). The model consists of two ODEs (one for each cell type), that each take the following general form:

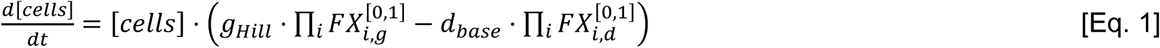

In Eq. 1, [*cells*] represents the density of that cell type in culture, *g_Hill_* represents density-inhibited cell proliferation using a Hill-like function, and *d_base_* represents cell death as a first-order reaction. In Eq. 1, there is a *FX_i_* function for each perturbant used in experiments, including: the progestin medroxy-progesterone acetate (MPA), the progesterone receptor inhibitor mifepristone (RU-486), the cytokine interleukin 1*β* (IL-1*β*), and the presence of other cell types in co-culture. The FX terms represent the effect of each perturbant on proliferation (*FX_i,g_*) or on cell death (*FX_i,d_*). These are on/off functions that either have no effect (*FX_i_^0^* = 1) or some constant effect (*FX_i_^1^* ≠ 1) depending on whether the perturbant is present. Once optimized, the magnitude of the FX_i_ terms represents whether the perturbant stimulates (*FX_i_* > 1), inhibits (0 < *FX_i_* < 1), or has no effect on (*FX_i_* = 1) proliferation or death. Estrogen is not included as an FX term because all experiments conducted in (22) include estradiol throughout the experimental timeframe (except for the withdrawal condition described below); hence, *g_Hil_* and *d_base_* represent the base proliferation and death of cells in the presence of estrogen.

The exact model equations and parameters used to predict the changes in the density of EEOCs (Eq. 2), and ESCs (Eq. 3) are as follows:

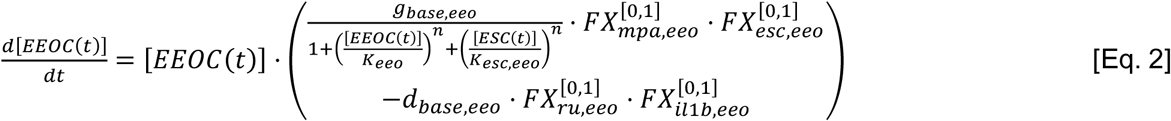

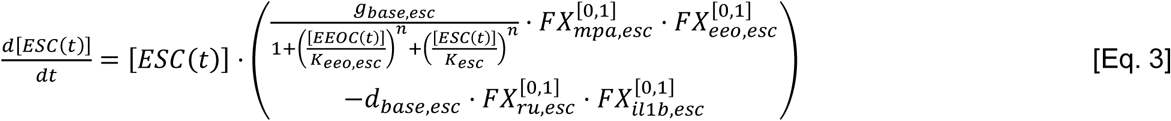

Contact inhibition of cell proliferation can be present in 2D and 3D cultures (40) and is the process by which cells limit entry into the cell cycle as the density of cells increases (41). To avoid modeling the complex mechanisms contributing to cell proliferation and its contact inhibition, we use a Hill-like function. Hill functions are commonly used to model systems where the rate of a process increases nonlinearly as the value of a variable within the system also increases until reaching a saturation point where the rate is maximized (42). For modeling processes that are inversely related to other variables, researchers alter the Hill function (32), producing a Hill-like equation of similar form to the Hill-like terms Eq. 2-3. In Eq. 2-3, the base cell proliferation rate (*g_base_*) is limited by the increasing density of cells (both EEOCs and ESCs). This cell-density-inhibited proliferation is tuned by Hill-like coefficients (*n*) and half-saturation constants (*K*) that represent how much each cell type inhibits their own proliferation (*K_eeo_*; *K_esc_*) and each other’s proliferation (*K_eeo,esc_*; *K_esc,eeo_*). Since epithelia and stroma can stimulate each other’s proliferation (17), the model also includes separate FX terms (*FX_eeo,esc_*; *FX_esc,eeo_*) as multipliers to *g_base_*; this allows for epithelia and stroma to have a positive effect on each other’s proliferation when they are present.

The FX terms representing progestin’s effect on each cell type (*FX_mpa,eeo_*; *FX_mpa,esc_*) are multipliers to the proliferation term because progestin is thought to inhibit the proliferation of estrogen-primed cells (39,43,44). During the menstrual cycle, menses occurs as the estrogen and progesterone levels decline; cell death in human endometrial cells is increased in response to estrogen and progesterone decline *in vitro* (45) and in late secretory/menses-staged tissues *ex vivo* (46,47). Hence, the FX terms representing RU-486 treatment (*FX_ru,eeo_*; *FX_ru,esc_*) are multipliers to the cell death term in the equations. Since IL-1*β* is also known to induce apoptosis in endometrial cells (22), its FX terms (*FX_il1b,eeo_*; *FX_il1b,esc_*) are multipliers to the cell death term in the equations.

### Simulating Cell Proliferation during Experiments

In the experimental study from which we draw data for optimization (22), an endometrial biopsy is taken from a single donor and then used to generate 15 parallel experimental conditions as outlined below. These experiments were repeated in their entirety using the biopsy-derived cells from 3 donors. These donors were of different ages (27 – 42 years old) and received different clinical diagnoses (endometriosis, fibroids, abnormal uterine bleeding), and the donors were in different phases of their menstrual cycles when samples were collected (Table S1). To identify optimal model parameter values for each donor, we simulated the 15 experimental culture conditions and compared our simulations to the experimental measurements (Fig. 1C).

The experimental results used for model optimization were obtained from 3D cell cultures in which cells were encapsulated within a hydrogel matrix (specifically, a polyethylene glycol (PEG)-based gel) which was surrounded by liquid culture media (Fig. 1B) and cultured for up to 15 days. The experiments investigated three different scenarios of cells included in the hydrogel (cell cases), which were each tested with five variations of hormones/cytokines present in the media (perturbant cases) (Fig. 1C). The three cell cases were:

- Monoculture experiment with EEOs;
- Monoculture experiment with ESCs;
- Co-culture experiment with both EEOs and ESCs.

The 5 media variations (perturbant cases) were:

- “Estrogen”: Estradiol added to the media throughout the experiment to mimic the proliferative phase of the menstrual cycle;
- “Estrogen, Progestin”: Estradiol and a progestin (MPA) added to the media throughout the experiment to mimic the secretory phase of the menstrual cycle;
- “Estrogen, Progestin, Withdrawal”: Estradiol and MPA added to the media (mimicking the secretory phase of the menstrual cycle) then removed 3 days prior to the end of the experiment, with the progesterone receptor inhibitor RU-486 added for the final 3 days to mimic menses;
- “Estrogen, IL-1*β*”: Estradiol and the cytokine IL-1*β* added to the media throughout the experiment, representing an inflamed proliferative phase;
- “Estrogen, progestin, IL-1*β*”: Estradiol, MPA, and IL-1*β* added to the media throughout the experiment, representing an inflamed secretory phase.

The *in vitro* experiments each lasted 12-15 days. We used Eq. 2-3 to simulate these experiments *in silico*, producing predictions for cell density and organoid growth during these experiments. To identify values for the model parameters, we used optimization approaches that compared our computational model predictions/simulations to the experimental data available. The experimental data included measurements of the size (diameter) of ∼11 individual EEOs (median) at each experimental timepoint and estimates of the relative density of ESCs at the end of each experiment based on day 15 Live/Dead staining (Fig. 1D-E). The computational model (Eq. 2-3) predicts cell density, the number of cells per volume of the encapsulating hydrogel, directly. To enable comparison with experimental data, we converted the simulated epithelial cell densities (*[EEOC]_i,j_*) to estimated values of average outer organoid radius (*R_i,j_*) at each time point (*j*) of each experiment (*i*) using Eq. 4-5:

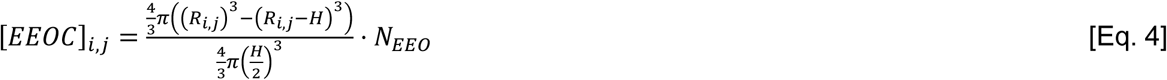

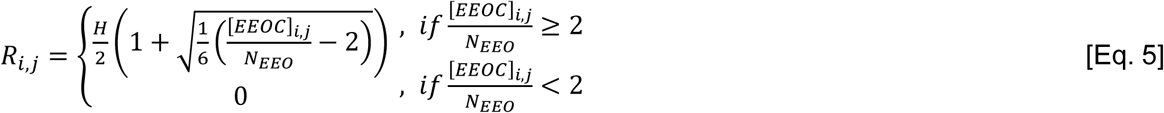

In Eq. 4, *[EEOC]_i,j_* is the number of epithelial organoid cells per volume of hydrogel (referred to as the “EEOC density”), *R_i,j_* is the average outer organoid radius, and *H* is the height of an individual epithelial cell within the organoid. *N_EEO_* represents the number of organoids per volume of hydrogel, which does not change over time or between experiments. Since an organoid is geometrically like a hollow sphere, we modeled the cell-containing volume of the organoid as a shell having an outer radius *R_i,j_* and a concentric inner radius of *(R_i,j_* – *H)*. We divided the cell-containing volume of the organoid by the estimated volume of one epithelial cell to obtain an approximate number of cells per organoid. We then multiplied the number of cells per organoid by the number of organoids per volume of hydrogel to yield the EEOC density (Eq. 4). The outer organoid radius is calculated from the cell density through rearranging Eq. 4 to solve for *R* in terms of *[EEOC]_i,j_* (Eq. 5).

The macroscopic nature and semi-spherical shape of EEOs makes it possible to measure their outer radius over time, from which cell density could be estimated. We use EEO measurements collected from the monoculture (EEO) and co-culture experiments that were conducted using cultures generated from each donor. Given we only have day-15 staining data for ESCs, and not time-course data, we normalize the predicted day-15 cell densities from each ESC monoculture experiment to the values for the estrogen-only case as shown in Eq. 6; the experimental data is similarly normalized.

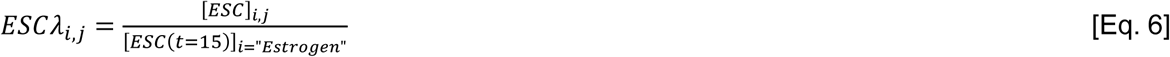

In Eq. 6, *ESCλ_i,j_* represents the relative ESC density for each time point and experiment, *[ESC]_i,j_* represents the stromal cell density for each time point and experiment, and *[ESC(t=15)]_i=”Estrogen”_* is the stromal cell density on day 15 of the estrogen-only monoculture experiment. The initial conditions used for our model simulations are defined in Table 1.

**Table 1.**
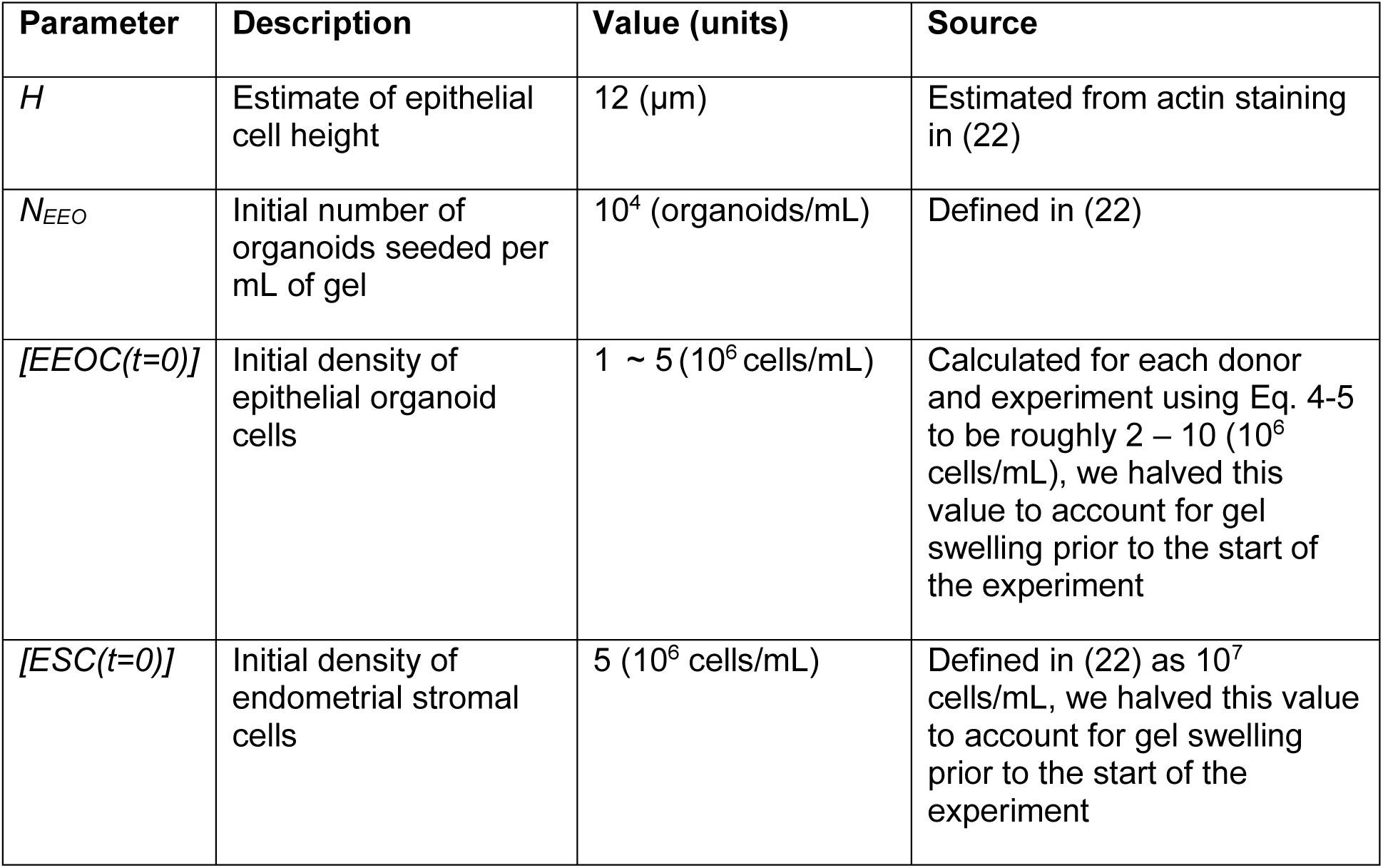
Initial conditions for simulating cell culture experiments that were conducted in (22)

### Optimizing the ODE Model Parameters and Quantifying Uncertainty

We used a non-linear least squares solver in Matlab (lsqnonlin) to identify a parameter set (“solution”) that minimizes the sum-squared error (*SSE*) between the measurements from experiments using a single donor’s cells and our model predictions (Eq. 7).

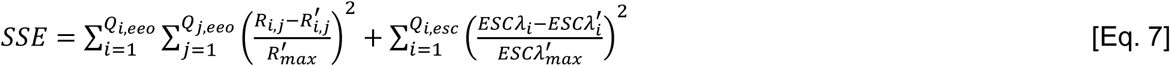

In Eq. 7, the experimental data for EEO radius and relative ESC density are indicated by *R′_i,j_* and *ESCλ′_i_* respectively, and our computational model predictions are represented by *R_i,j_* and *ESCλ_i_*. Since multiple organoids were measured at each time point, *R′_i,j,_* represents the mean organoid radius at each time point. The number of data points per experiment (*Q_j_*) and the number of experiments with data collected (*Q_i_*) can vary by donor, cell type, and condition. Experiments in (22) discern relative ESC density from the monoculture day-15 Live/Dead staining. Hence, *ESCλ′_i_* refers to the Live/Dead staining measurement for each perturbation case normalized to the measurement for the estrogen-only monoculture case. *R′_max_* and *ESCλ′_max_* represent the maximum outer organoid radius and day-15 relative ESC density from the experimental measurements, respectively. The values for parameters used in optimization were defined to match the experimental design of (22) and are provided in Table 2.

**Table 2.**
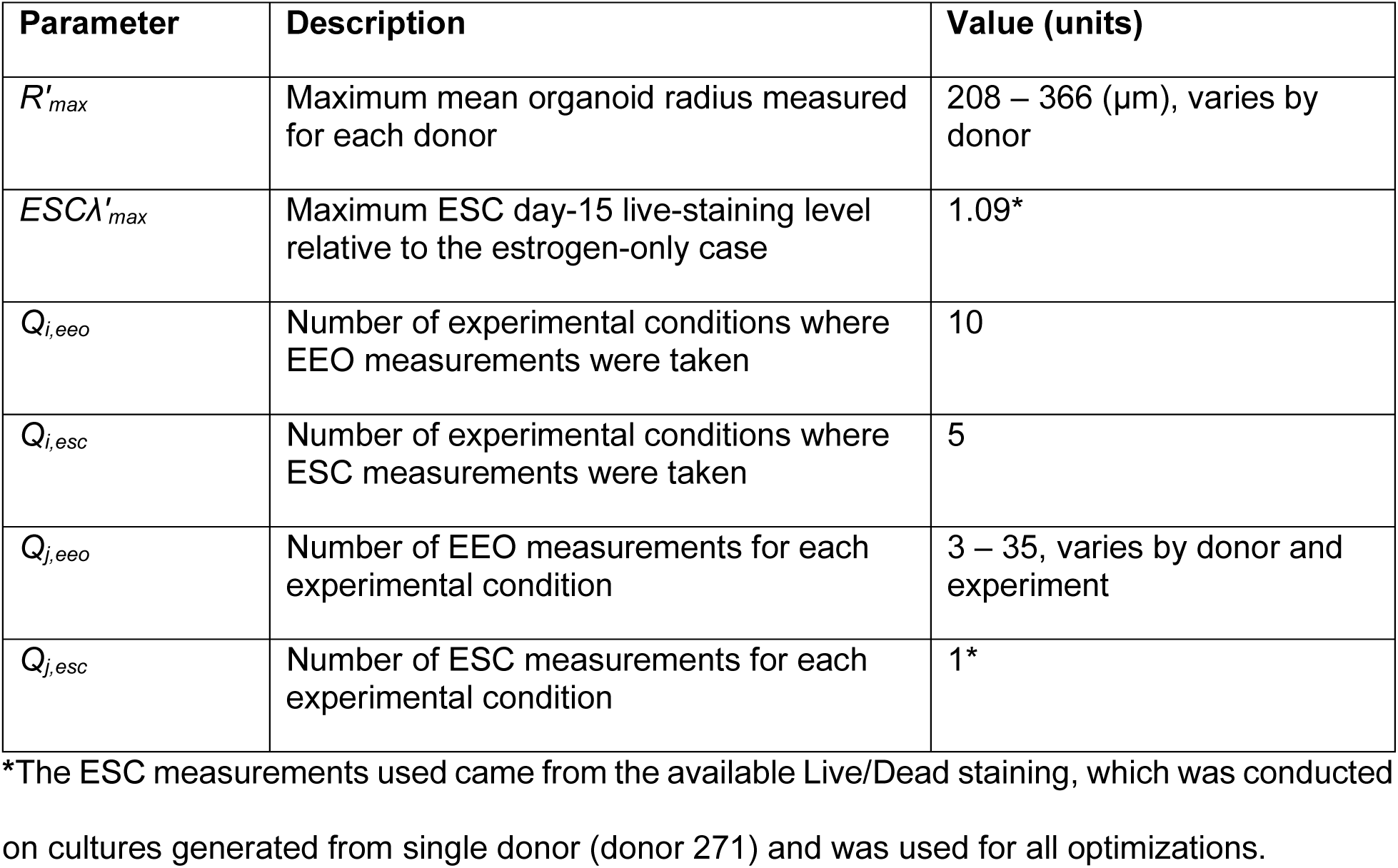
Parameters used in optimization sum-squared error calculation (Eq. 7)

To identify uncertainty in our optimized parameter values, we ran lsqnonlin 100 times in total, initializing it each time at a new starting estimate for model parameter values within our defined bounds (Table 3). We then gated the resulting parameter sets to only include those with predictions similar to experiments. Parameter sets that were included in our estimates of uncertainty must not have:

- **“Excess error”**, specifically that the model predictions must be within +/- 65% of the corresponding measured organoid means at ≥90% of the experimental time points. This threshold was set to match the uncertainty in the experimental data (Fig. S1). Although we use the mean organoid measurements for each experiment involving EEOs in optimization, these means were calculated from measurements of individual organoids that could deviate by as much as 3.6X (i.e., 260% more than) the calculated mean. Since 90% of the experimental measurements fell within +/- 65% of the calculated mean (Fig. S1), we defined this as a quality threshold for our optimizations.
- **“Excess epithelial growth”**, specifically that the simulated epithelial organoid radius must be plateauing or decreasing by day 18. Meaning, for the 10 mono- and co-culture EEO experiments, EEO cell density cannot be predicted to have both a positive first and second derivative after day 18 of simulation. This limit was set to prevent cases of unbounded proliferation and organoid growth, which was not observed experimentally.
- **“Excess stromal growth/death”**, meaning the ratio between the simulated day-15 and day-0 relative ESC density (*ESCλ_i,j_* in Eq. 6) must be less than 10^7^ but greater than 0.5. This limit was set to ensure that the growth of stroma does not exceed ∼10 million times the initial cell density. This upper limit corresponds to exponential growth with a doubling time of 18 hours (net growth rate = 0.92 day^-1^). For reference, 60 commonly used cancer cell lines have a doubling time ranging from 17.4 – 79.5 hours (48). We expect these immortalized cell lines to proliferate faster than the primary cells we are studying. This limit also ensures that the death of stroma does not exceed half the initial cell density since staining in (22) showed stroma in abundance on day 15 of monoculture.

**Table 3.** Bounds for calibration of parameters in the endometrial proliferation model (Eq. 2-3)

| Parameter(s) | Description | Range (units) |
| --- | --- | --- |
| $g_{base,eeo}$<br>$g_{base,esc}$ | Base proliferation rate constants for endometrial epithelial organoid cells (EEOCs) and endometrial stromal cells (ESCs) | $10^{-4} - 2$ (1/days) |
| $d_{base,eeo}$<br>$d_{base,esc}$ | Base death rate constants for EEOCs and ESCs | $10^{-3} - 1$ (1/days) |
| $K_{eeo}$<br>$K_{esc}$<br>$K_{eeo,esc}$<br>$K_{esc,eeo}$ | Half-saturation rate constant for each cell type's proliferation | $1 - 2000$<br>($10^6$ cells/mL) |
| $n$ | Hill-like coefficient for the effect of cells on proliferation | $0.1 - 10$ (unitless) |
| $FX_{eeo,esc}$<br>$FX_{esc,eeo}$ | Constants modeling the stimulatory or inhibitory effect of EEOCs on ESCs and vice versa | $1 - 10^2$ (unitless) |
| $FX$ terms<br>(all others) | Constants modeling the stimulatory or inhibitory effect of adding perturbant to media on each cell type's proliferation | $10^{-2} - 10^2$ (unitless) |

Having gated the parameters as stated above, we calculated and plotted summary statistics for each parameter. We also identified the **“best solution”**, which we defined as the parameter set that produced simulations with the lowest SSE (Eq. 7) out of the parameter sets that were accepted by the above criteria.

### Simulating IL-1*β* Reaction-Diffusion in 3D Hydrogel Cell Cultures Using PDEs

To complement the ODE-based computational model of cell proliferation, we created a PDE-based reaction-diffusion model to predict the spatial distribution of IL-1*β* after it’s been added exogenously in the liquid media surrounding the 3D hydrogel cell cultures generated in (22). Diffusion is modeled according to Fick’s second law with a reaction term (Eq. 8).

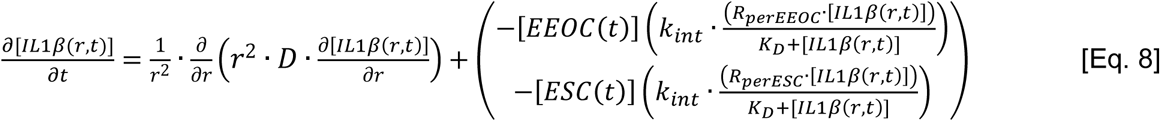

In Eq. 8, [*IL1β(r,t)*] represents the concentration of IL-1*β* along the radius *r* of the hemispherical hydrogel over time *t*, with the radius being that of the final gel volume after the hydrogel swells in media prior to t=0. *D* is the diffusion coefficient for IL-1*β* within a hydrogel containing cells and *k_int_* is the rate constant for IL-1*β* consumption via cell-surface receptors. Using a pseudo-steady-state assumption, the quantity of IL-1*β*:Receptor complexes was approximated as the ratio between the total available receptors per cell (*R_perEEOC_* and *R_perESC_*), the concentration of IL-1*β*, and the equilibrium dissociation constant for receptor-binding (*K_D_*).

We used our model of epithelial and stromal cell density (*[EEOC(t)]* and *[ESC(t)]*) (Eq. 2-3) to calculate changes in receptor densities, assuming that cells are homogenously distributed throughout the hydrogel. We solved the reaction-diffusion equation along the radius of these hemispherical hydrogels in spherical coordinates using Matlab, with boundaries set at the center of the hydrogel (*r*=0) and the gel edge (*r*=*r_gel_*). We assumed angular symmetry, constant concentration in the media at the gel edge, and no flux through the gel center. The model was simulated at equilibrium swelling conditions, meaning the gel radii used in simulations were for the final volumes after gel swelling in media with no net flux of aqueous media into or out of the gel due to swelling. Since the PEG-based gel used in (22) could double in size after swelling (21,49,50), we calculated the gel radius using double the initial gel volume for these cultures. The model parameters for simulating IL-1*β* diffusion and reaction are summarized in Table 4 and derived in the *Supplemental Methods*.

**Table 4.** Parameter values used to simulate IL-1*β* reaction-diffusion in the 3D culture system that was used to calibrate this computational model (22). Calculations are in *Supplemental Methods*.

|  | Parameter | Value (units) | Estimated from | Range explored here |
| --- | --- | --- | --- | --- |
| $r_{gel}$ | Gel radius (post-swelling to double the initial gel volume) | 0.142 (cm) | (22) | 0.06 – 0.3 cm |
| $V_{gel}$ | Gel volume (post-swelling) | 6 (μL) | (22) | 0.6 – 60 |
| $u_{IEM}$ | IL-1β maintained in media | 0.06 (nM)<br>1 (ng/mL)<br>$3.5 \times 10^{10}$ (#/mL) | (22) | n/a |
| $K_D$ | IL-1β-receptor binding affinity | 2.2 (nM)<br>37.4 (ng/mL)<br>$1.3 \times 10^{12}$ (#/mL) | (51) | n/a |
| $D$ | Diffusion coefficient for IL-1β through gel | $5 \times 10^{-7}$ (cm<sup>2</sup>/s) | (52) | $5 \times 10^{-8}$ – $5 \times 10^{-6}$ |
| $k_{int}$ | Receptor internalization rate | $10^{-4}$ (1/s) | (53) | $10^{-5}$ – $10^{-3}$ |
| $R_{percm2}$ | Receptors per cell surface area | $8.5 \times 10^8$ (#/cm<sup>2</sup>) | (54) | $8.5 \times 10^7$ – $8.5 \times 10^9$ |
| $R_{perEEOC}$ | Total accessible receptors on EEOCs | 800 (#/cell) | (54) | 80 – 8000 |
| $R_{perESC}$ | Total accessible receptors on ESCs that are accessible for binding | 1700 (#/cell) | (54) | 170 – 17000 |

## Results

We first present results from optimizing and analyzing our computational model of proliferating endometrial cells in 3D culture, then we apply the results of this model to consider the effects of cell proliferation on molecular gradient formation within these culture systems.

### The ODE-based computational model of proliferating endometrial cells was parametrized using data from mono- and co-culture experiments

To identify sets of parameter values that define the rates of proliferation and death of endometrial cells, we used our model, optimization algorithms, and experimental measurements of organoid radius from time-lapse imaging and of day-15 relative ESC density from Live/Dead staining. To assess the uncertainty in the parameter values, we initialized our optimization solver from 100 different starting points and then applied gating to only include plausible solutions, as described in the *Methods*.

We used this method to identify donor-specific parameter values that resulted in simulation predictions that matched the donor-specific experimental data. The optimization for donor 272 achieved the lowest sum-squared error overall. Out of 100 optimizations, 81 were acceptable by our gating criteria and had similar predictions for EEO growth across our simulations of the *in vitro* experiments. Out of the remaining 19 solutions, 9 solutions were removed because of excess epithelial growth, 7 solutions were removed due to excess stromal growth or death, 2 solutions were removed due to failing both the epithelial and stromal gates, and 1 solution was removed due to excess error (Fig. 2, Fig. S2). The accepted parameter set with the lowest sum-squared error simulated the results of all 15 experiments well, with model predictions for EEOs falling within 20% of the experimental measurement means at 98% of the time points. The largest source of variability was in the predictions for ESC density in co-culture, which is to be expected as data for ESCs in co-culture was limited.

**Figure 2.**
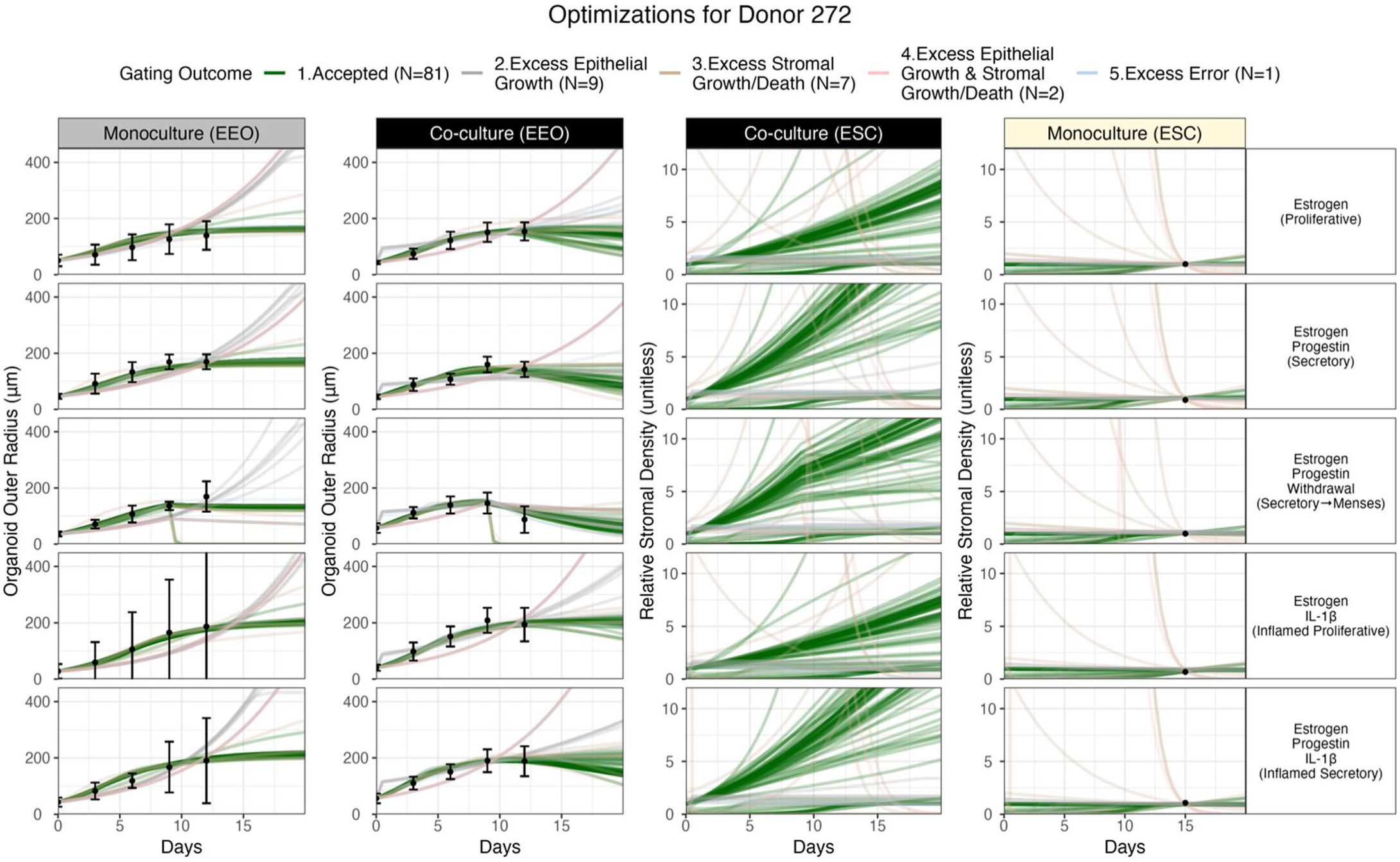
Optimization of the computational model to experimental data gives a set of non-unique but constrained parameter set solutions (example donor: 272). Each row represents one perturbation condition (noted on right-hand side), and the columns represent the culture condition and cell type (noted at top); thus the twenty panels represent the outcomes of 15 experiments (dots) and their equivalent 15 simulation conditions (lines). Each line represents a simulation with a different parameter set, each parameter set obtained by optimization to the experimental data. We ran 100 optimizations to identify consistent parameter sets for cells within endometrial epithelial organoids (EEO) and endometrial stromal cells (ESC) from a donor with abnormal uterine bleeding. Each optimized parameter set produced simulations for all 15 experimental conditions. The resulting simulations are color-coded based on our gating analysis to indicate whether they were accepted as solutions for further analysis (green) or removed for one of the following reasons: excess error (blue), excess epithelial growth (gray), excess stromal growth/death (brown), or a combination of both excess epithelial growth and excess stromal growth/death (red), as described in *Methods*. For experimental data, the black circles represent the mean EEO measurements and the relative ESC density from day-15 Live/Dead-staining data that were used during optimization, and the error bars represent the standard error in EEO measurements. The corresponding parameter sets are shown in Supplemental Figure S2, and the equivalent figures for other donors are in Supplemental Figures S4 and S5.

We used the same optimization method to estimate model parameters for donor 271 (Fig. S4) and donor 260 (Fig. S5). For these donors, there were more solutions removed due to excessive epithelial growth, but in each case there remained 33 and 18 accepted parameter sets, respectively (Fig. S4-S5). There was also consistency in the accepted vs rejected solutions; for example, looking at the parameter distributions for donor 260, the solutions removed due to excess epithelial growth showed IL-1*β* inhibiting stromal cell death (Fig. S5B: values of *FX_il1b,esc_* below 1) whereas the corresponding term for the accepted cases stimulated stromal cell death (values of *FX_il1b,esc_* above 1).

Although the simulations and experimental data are shown in Figure 2 in the units measured experimentally (outer organoid radius, relative stromal cell density), we converted these units to absolute cell density for simulations (using Eq. 4 & 6) in order to compare the simulations and parameter values for EEOs and ESCs on the same scale (Fig. 3–4, Fig. S6). Using Eq. 4. and the initial conditions (Table 1), an EEO outer radius of 100 µm converts to ∼10^7^ cells/mL. A relative ESC density of 1 translates to ∼10^7^ cells/mL in our best solution for each donor (Fig. S6).

**Figure 3.**
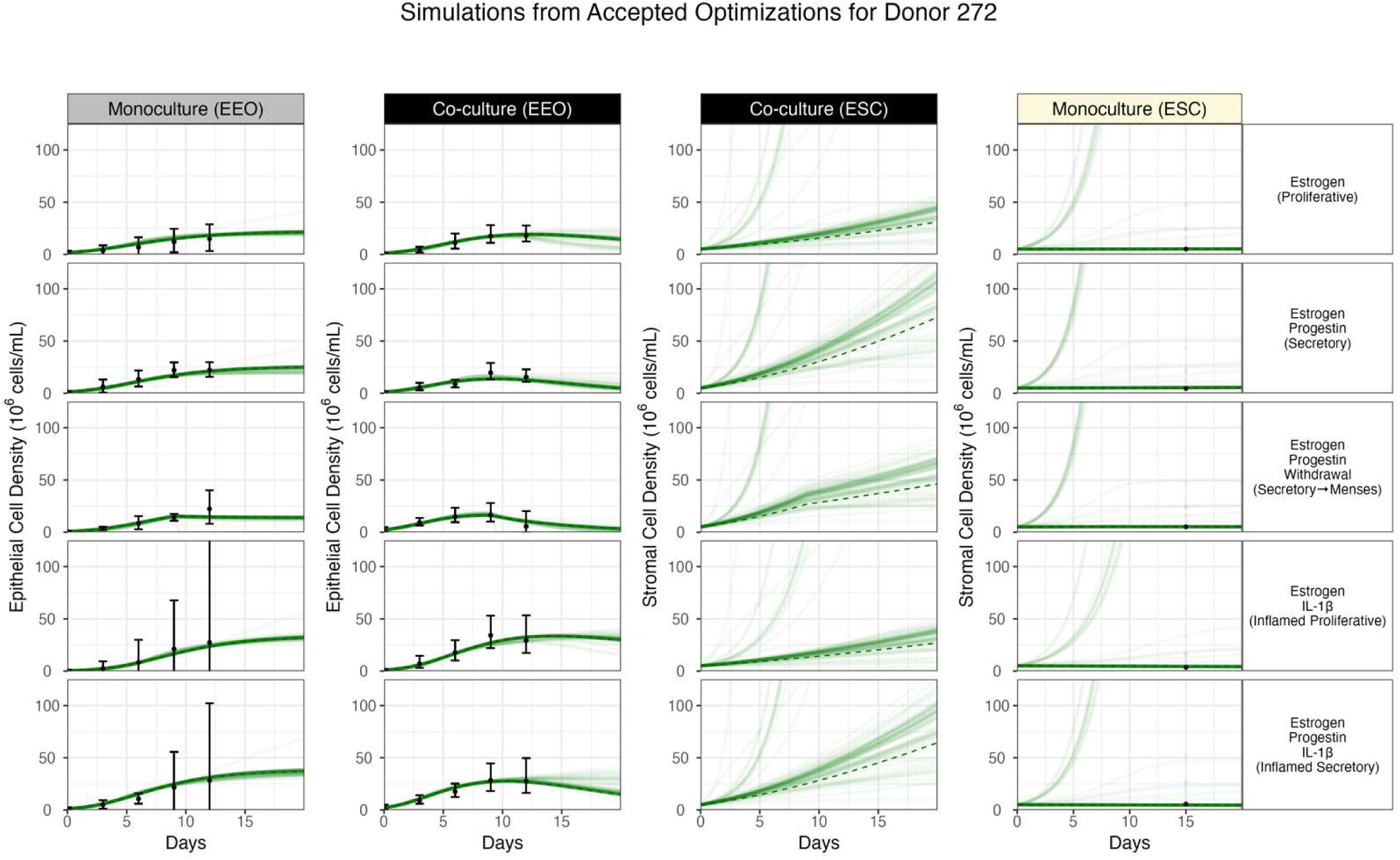
Uncertainty in optimized model simulations for accepted parameter sets (example donor: 272). This figure presents the accepted portion of model optimizations from Figure 2 converted to absolute units of cell density for both epithelial and stromal cells (as described in *Methods*). The black dots represent the mean organoid experimental measurements converted to cell density and the relative ESC density converted to absolute cell density for each optimization result. Simulations from the best solution are denoted with a dashed line. The “best” solution is the parameter set that produced simulations with the lowest sum-squared error (Eq. 7) out of the parameter sets that were accepted according to the criteria described in *Methods*.

**Figure 4.**
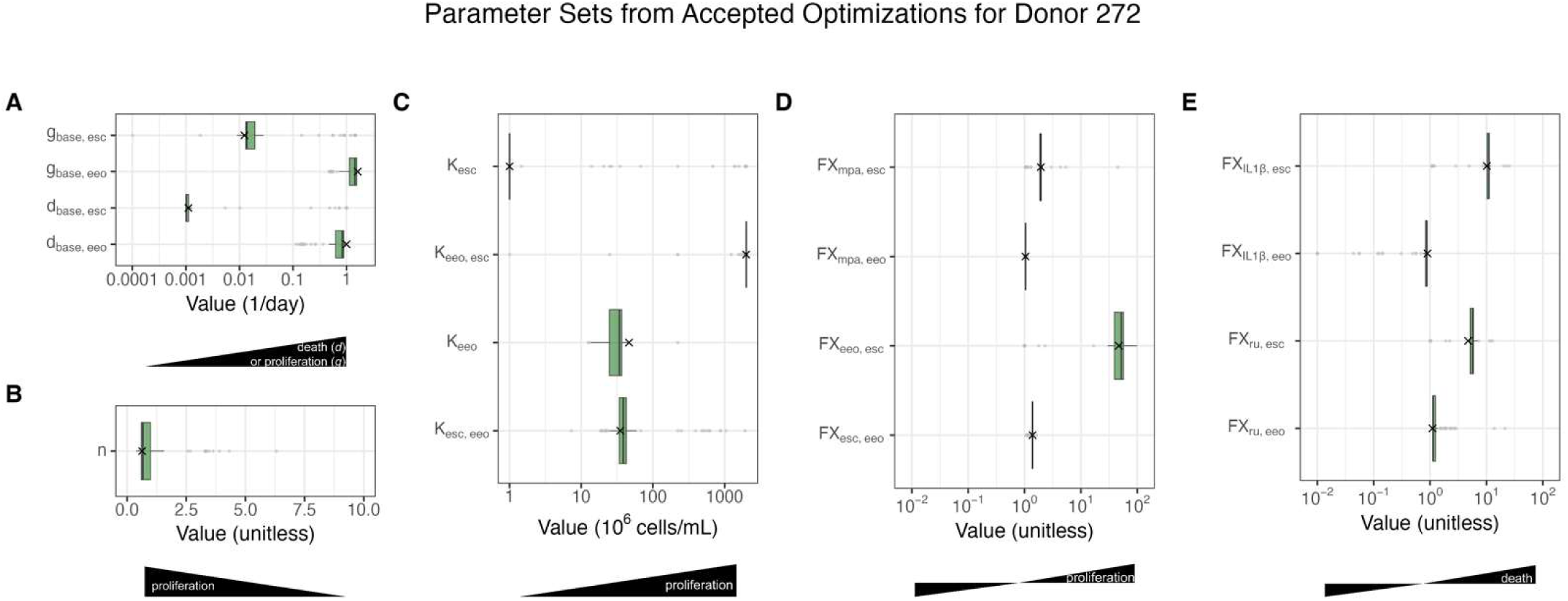
Optimized model parameters for donor 272. Distributions of the values of each parameter in the accepted solutions (i.e. after gating). Parameter values for the best solution are denoted with X’s and the numerical values are listed in Table 3. The “best” solution is the parameter set that produced simulations with the lowest sum-squared error (Eq. 7) out of the parameter sets that were accepted according to the criteria described in *Methods*. Parameters are arranged by role in increasing or decreasing the net growth rate (where net growth rate = proliferation rate – death rate): **(A)** Base proliferation and death rates, **(B)** Hill-like coefficient for proliferation, **(C)** half-saturation rate constant for proliferation, **(D)** perturbant effects on proliferation, and **(E)** perturbant effects on cell death. The black annotations below the x-axis aid in interpreting the parameter values by showing the direction in which cell proliferation or death increases or decreases. Key for each box plot: middle bar = median; lower/upper bar = 25^th^/75^th^ percentile; whiskers = least and greatest values (no further than 1.5 ⨯ the inter-quartile range); gray circles = outliers. Narrow ranges of accepted parameters (I.e. narrow box plots) indicate well-constrained parameters.

Optimization and gating allowed us to identify for each donor: (i) multiple sets of parameters that could recreate all experiment results and (ii) a measure of the uncertainty in both model simulations and parameter values. Comparing the simulations for the best solutions that passed gating, we see that the model simulations match the experimental data and predict the dynamics of quantities not measured, such as stromal cell density (Fig. 5). Next, we use our model to compare epithelial growth and predicted stromal cell density in mono- and co-cultures from different donors.

**Figure 5.**
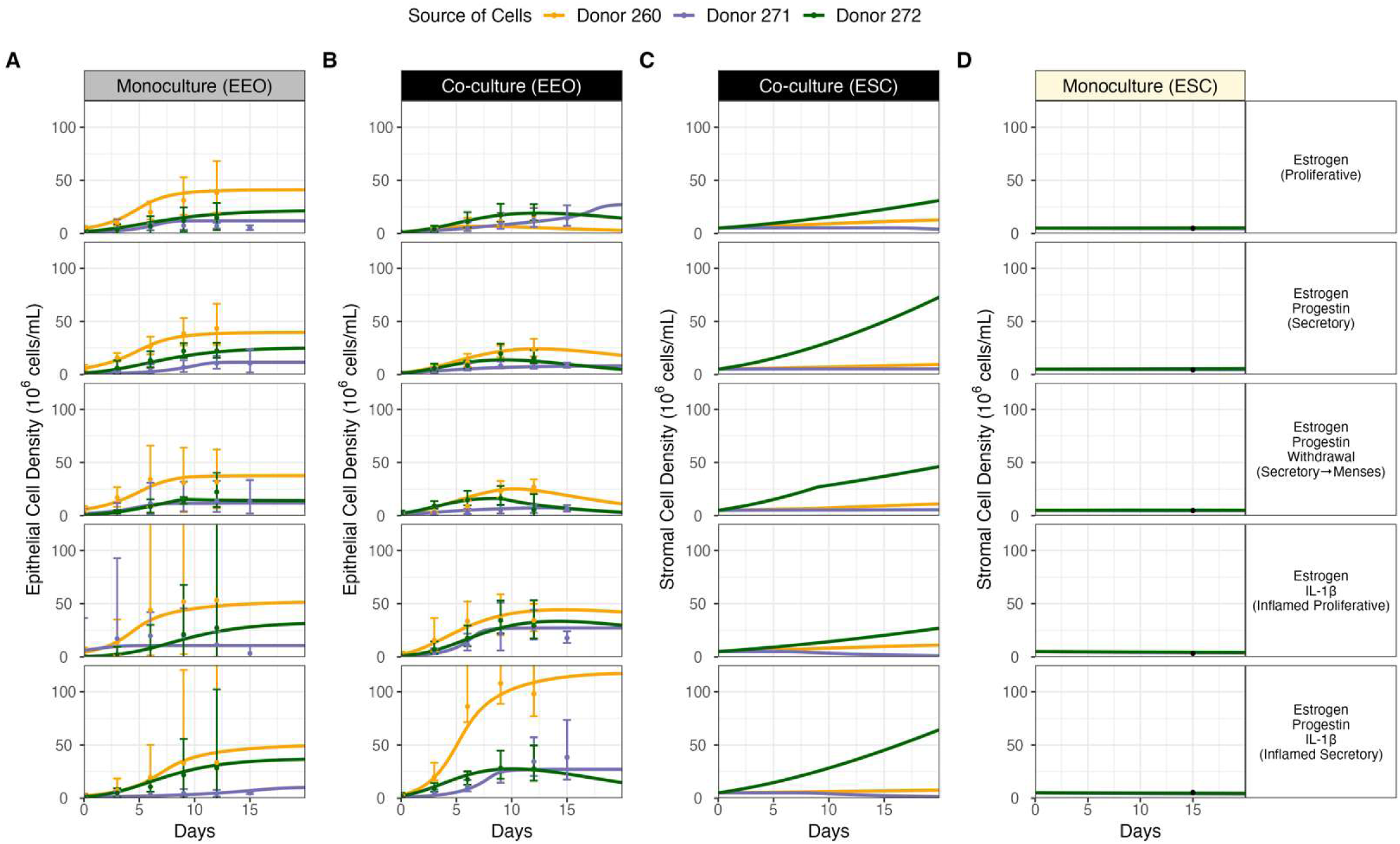
Comparison of optimized model simulations and experimental results across donors. Simulation results from the best-fit accepted parameter sets for each donor (as described in *Methods*), including: donor 272 (shown previously, green), donor 271 (purple), and donor 260 (orange). Simulations (lines) and experimental data (dots) are plotted in terms of cell density for the 5 perturbation cases (rows) conducted in 3 cell culture conditions: **(A)** monoculture with EEOs, **(B-C)** co-culture of EEOs and ESCs, and **(D)** monoculture with ESCs. The dots represent the mean organoid experimental measurements converted to cell density using Eq. 4.

### Endometrial epithelial cells may outgrow stromal cells in monoculture and have higher turnover (faster proliferation, faster death) than stromal cells

For cells derived from each of the donors, our model predicted that the net increase in epithelial cell density in monoculture was greater than that of stromal cell density in the corresponding monoculture experiments (Fig. 5A, 5D); this was the case even with less stromal cell experimental data than epithelial data to constrain its simulations. Accordingly, our optimization found epithelial cells to have higher base proliferation rates (*g_base_*) than stromal cells across cultures generated from different donors (Fig. 6A; Table 5). The half-saturation rate constants for self-inhibition of growth (*K_eeo_* and *K_esc_*) were slightly higher for EEOs than ESCs (Table 5) from each donor, suggesting more growth inhibition (lower proliferation) for stromal cells at similar cell densities. Interestingly, the base death rates (*d_base_*) were also higher for the epithelial cells compared to stromal cells. The balance of proliferation and death leads to the observed differences in cell density and organoid size. Therefore, even with higher base death rates, net change in epithelial cells density in monoculture is higher than that of stromal cells, suggesting higher overall turnover of the epithelial cells than the stromal cells. The impact of the other terms in the equations (e.g. *FX* – see Table 5) influence this growth rate in the different experimental conditions.

**Figure 6.**
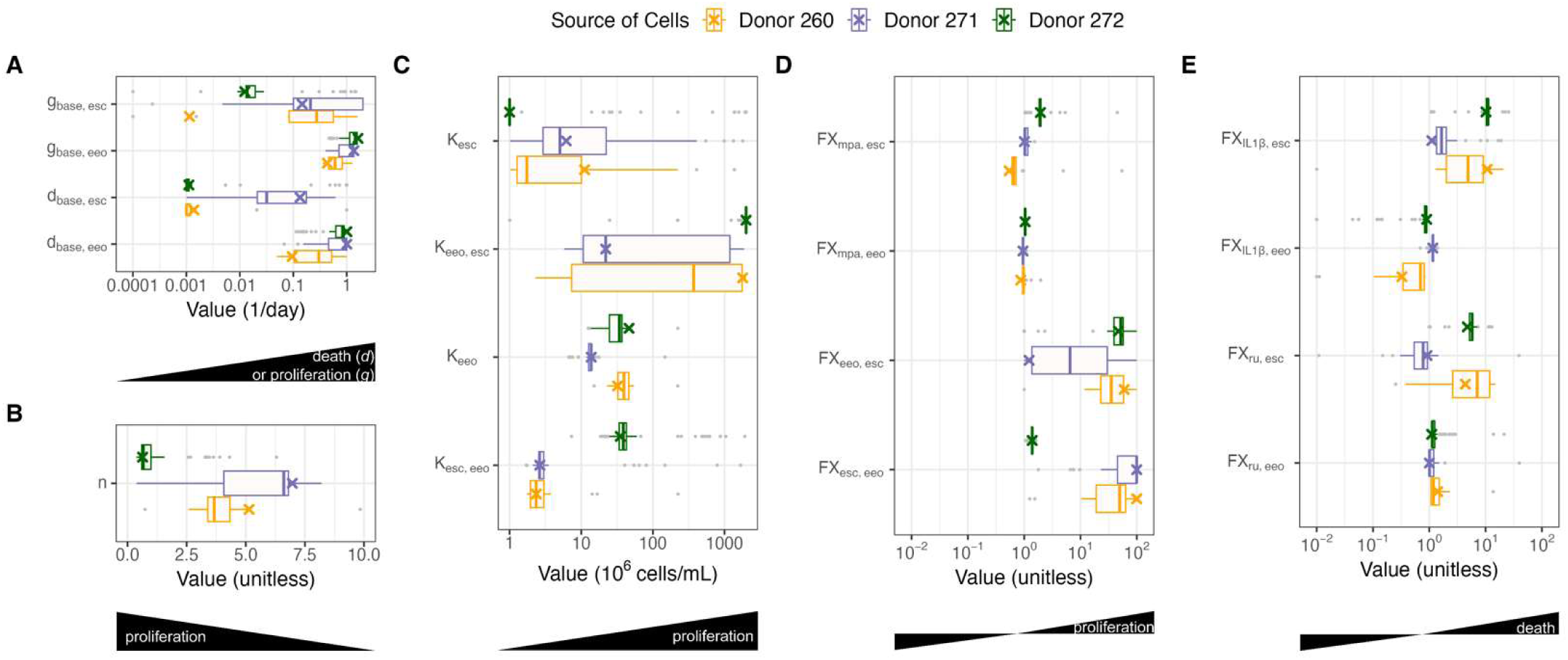
Optimized model parameter values compared across donors. Distribution of parameter values for the accepted parameter sets for each donor (as described in *Methods*). Parameter values for the best solution are denoted with X’s and the numerical values are listed in Table 5. The “best” solution is the parameter set that produced simulations with the lowest sum-squared error (Eq. 7) out of the parameter sets that were accepted according to the criteria described in *Methods*. Parameters are arranged by role in increasing or decreasing the net growth rate (where net growth rate = proliferation rate – death rate): **(A)** Base proliferation and death rates, **(B)** Hill-like coefficient for proliferation, **(C)** half-saturation rate constant for proliferation, **(D)** perturbant effects on proliferation, and **(E)** perturbant effects on cell death. The black annotations below the x-axis aid in interpreting the parameter values by showing the direction in which cell proliferation or death increases or decreases. Key for each box plot: middle bar = median; lower/upper bar = 25^th^ /75^th^ percentile; whiskers = least and greatest values (no further than 1.5 ⨯ the inter-quartile range); gray circles = outliers.

**Table 5.**
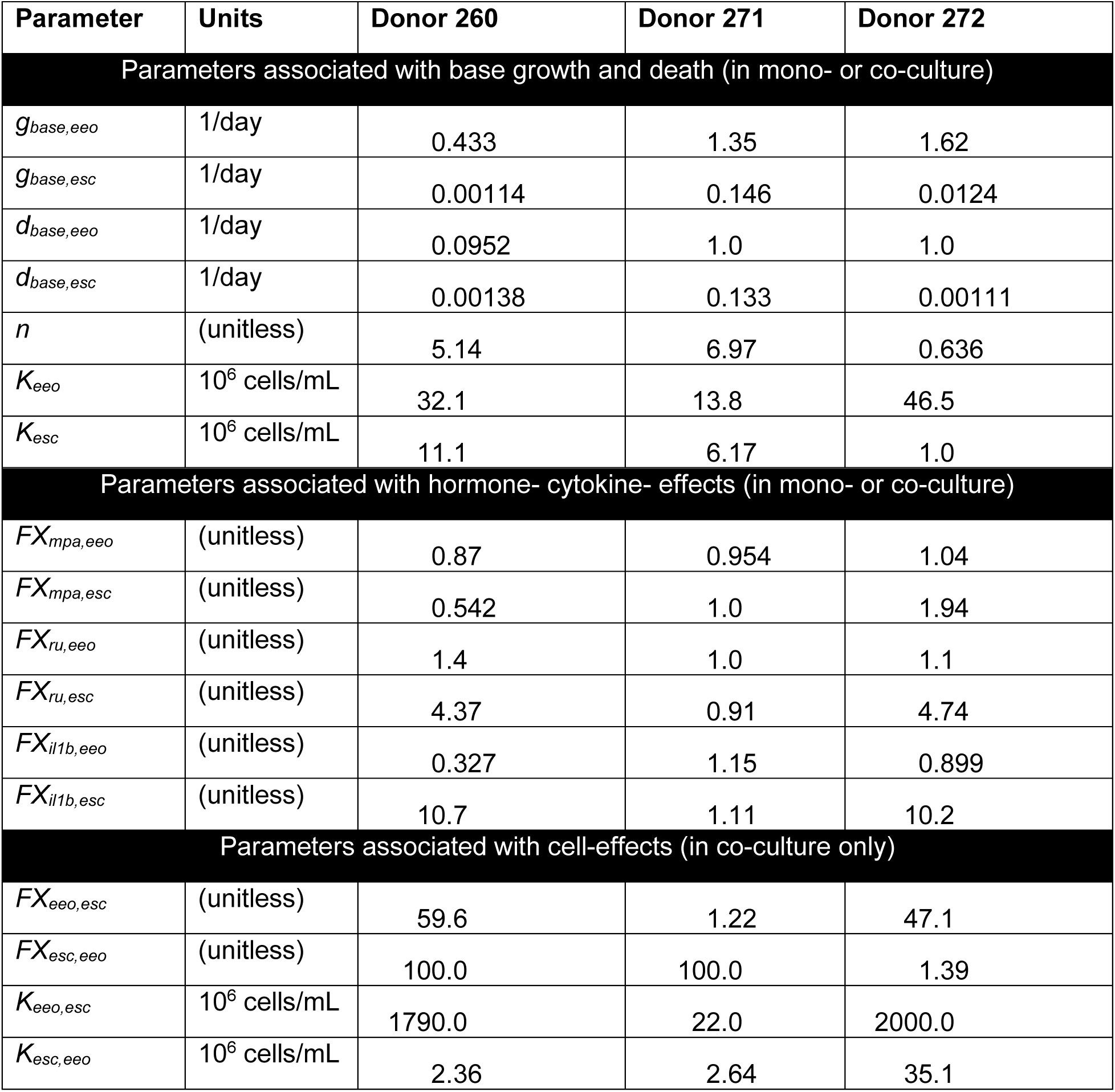
Optimized parameters for cultures derived from each donor for the best solution (the solution with lowest sum-squared error out of those accepted following gating, as described in the *Methods*). Parameter value distributions for all accepted solutions are shown in Figure 6.

| Parameter | Units | Donor 260 | Donor 271 | Donor 272 |
| --- | --- | --- | --- | --- |
| Parameters associated with base growth and death (in mono- or co-culture) |  |  |  |  |
| $g_{base,eeo}$ | 1/day | 0.433 | 1.35 | 1.62 |
| $g_{base,esc}$ | 1/day | 0.00114 | 0.146 | 0.0124 |
| $d_{base,eeo}$ | 1/day | 0.0952 | 1.0 | 1.0 |
| $d_{base,esc}$ | 1/day | 0.00138 | 0.133 | 0.00111 |
| $n$ | (unitless) | 5.14 | 6.97 | 0.636 |
| $K_{eeo}$ | $10^6$ cells/mL | 32.1 | 13.8 | 46.5 |
| $K_{esc}$ | $10^6$ cells/mL | 11.1 | 6.17 | 1.0 |
| Parameters associated with hormone- cytokine- effects (in mono- or co-culture) |  |  |  |  |
| $FX_{mpa,eeo}$ | (unitless) | 0.87 | 0.954 | 1.04 |
| $FX_{mpa,esc}$ | (unitless) | 0.542 | 1.0 | 1.94 |
| $FX_{ru,eeo}$ | (unitless) | 1.4 | 1.0 | 1.1 |
| $FX_{ru,esc}$ | (unitless) | 4.37 | 0.91 | 4.74 |
| $FX_{il1b,eeo}$ | (unitless) | 0.327 | 1.15 | 0.899 |
| $FX_{il1b,esc}$ | (unitless) | 10.7 | 1.11 | 10.2 |
| Parameters associated with cell-effects (in co-culture only) |  |  |  |  |
| $FX_{eeo,esc}$ | (unitless) | 59.6 | 1.22 | 47.1 |
| $FX_{esc,eeo}$ | (unitless) | 100.0 | 100.0 | 1.39 |
| $K_{eeo,esc}$ | $10^6$ cells/mL | 1790.0 | 22.0 | 2000.0 |
| $K_{esc,eeo}$ | $10^6$ cells/mL | 2.36 | 2.64 | 35.1 |

The effects of estrogen, progestin, and IL-1*β* on stromal cell density were smaller than their effects on epithelial cell density in monoculture (Fig. 5A, 5D). In order for these molecules to have visible effects on stromal cell density in monoculture simulations, cell proliferation would need to be high and cell death low without the hormone/cytokine present; this could be achieved through stromal cells having a high base proliferation rate constant (*g_base,esc_*), a low base death rate constant (*d_base,esc_*), and low self-inhibition (large *K_esc_*). Although the optimization algorithm had those conditions available to it, self-inhibition for stroma was consistently optimized to be higher than that of epithelia.

In contrast to monoculture (Fig. 5D), stromal cells in co-culture with epithelial organoids are predicted to show a greater net change in cell density (Fig. 5C, Fig. S3), and the direction (increase or decrease) differed for cells from the three donors. This suggests that stromal cell mitotic activity is heavily influenced by crosstalk with the epithelial cells. Similarly, though less dramatically, epithelial growth in co-culture (Fig. 5B) is different than in monoculture (Fig 5A), also reflecting the impact of epithelial-stromal cell crosstalk. These differences are well captured in the simulations, matching the experimental time-course measurements.

### Model simulations suggest that differences in cell responses to estrogen, progestin, and IL-1*β* can be explained by differences in epithelial-stromal regulation between donors

In both experiments and model simulations, the effects of hormones and IL-1*β* on cell density varied by the cell types and other molecules present in the media. To further quantify the effect of a hormone/cytokine on cell density, we calculated the ratio of the area under the curve (AUC) of the simulated cell density (Fig. 5) for one culture condition containing that hormone/cytokine to the AUC for another culture condition lacking that hormone/cytokine; these ratios are calculated for each donor (Fig. 7). Differences in AUC between culture conditions can be thought of as reflecting differences in “overall growth” or “overall change in cell density”.

**Figure 7.**
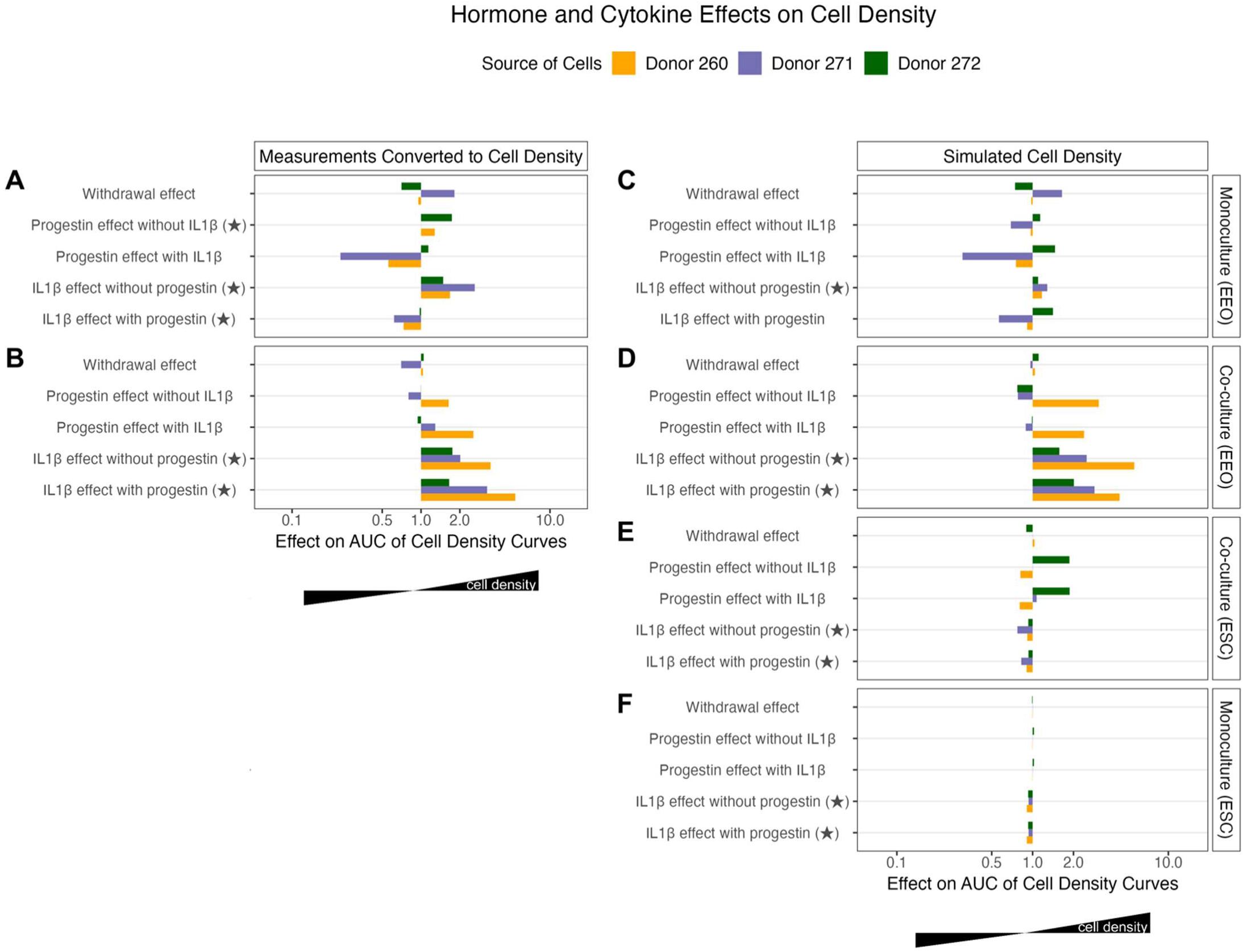
Quantifying specific perturbant effects on the cell densities across different culture conditions and source of cells. For each culture condition in Figure 5, we calculated the area under the curve (AUC) for the **(A-B)** experiment measurements (converted to units of cell density) and **(C-F)** the simulated cell densities produced from optimizing the computational model. The effect was then calculated as the ratio between the AUCs with and without the perturbant(s). More specifically: “Withdrawal effect” is the ratio between the estrogen-progestin-withdrawal condition and the estrogen-progestin condition; “progestin effect without IL1β” is the ratio between the estrogen-progestin condition and the estrogen condition; “progestin effect with IL1β” is the ratio between estrogen-progestin-IL1β condition and the estrogen-IL1β simulation; “IL1β effect without progestin” is the ratio between estrogen-IL1β condition and the estrogen condition; and “IL1β effect with progestin” is the ratio between the estrogen-progestin-IL1β condition and the estrogen-progestin simulation. The experiment measurements and simulations for each condition are shown in Figure 5. The star (★) denotes where perturbant(s) consistently increased or decreased the cell density across donors. The black annotations below the x-axis aid in interpreting the parameter values by showing the direction in which cell density would increases or decreases.

Some of the hormone/cytokine effects were consistent across cells from all three donors; most notably, the measured and simulated effect of IL-1*β* without progestin on the overall growth of EEOs (all strongly positive, Fig. 7B,D) and the simulated effect of IL-1*β* on the overall change in ESC density (all slightly negative, Fig. 7E-F). These effects of IL-1*β* were seen when comparing the “Estrogen, IL-1*β”* case to the “Estrogen” case (mimicking the effect of inflammation on the proliferative phase) and the “Estrogen, Progestin, IL-1*β”* case to the “Estrogen, Progestin” case (mimicking the effect of inflammation on the secretory phase and disruption of progesterone signaling). In experiments using different donor cells, the addition of progestin and IL-1*β* consistently increased the overall growth of EEOs more in co-culture than in monoculture (Fig. 7A-B), and our computational model was able to capture that difference (Fig. 7C-D). Other effects were donor-specific or experimental condition-specific. For example, in experiments using cells from donor 260, EEOs grew more in mono- and co-cultures in the secretory experiment than in the proliferative experiment (“progestin effect without IL1β” in Fig. 7A-B). When compared to measurements from the inflamed proliferative experiment, EEOs grew less in the inflamed secretory monoculture experiment; however, EEOs grew more in that co-culture experiment (“progestin effect with IL1β” in Fig. 7A-B). Of these four responses to progestin, our computational model captured well the responses to progestin in co-culture (both with and without IL-1*β*) and the response in monoculture with IL-1*β* (Fig. 7C-D). Since our optimized parameters estimated progestin to be slightly inhibitory of epithelial cells from donors 260 and 271 (*FX_mpa,eeo_* < 1) and epithelial self-inhibition to be low (*K_eeo_* > *[EEOC(t)]*) (Fig. 6C-D, Table 5), our model simulations showed less overall EEO growth in the secretory monoculture than in the proliferative monoculture for those two donors. However, the increase in this experiment measurement (Fig. 7A) and the decrease in the corresponding simulation prediction (Fig. 7C) were both small, and the simulations still fit the experimental data well (Fig. 5A).

Overall, our computational model reflects these between-experiment differences well because it includes separate parameters representing how epithelial and stromal cells affect each other’s proliferation (Eq. 2-3). For example, our optimization estimated parameters for the computational model for donor 260’s cells such that stroma slightly inhibited epithelial proliferation (*K_esc,eeo_* < *[ESC(t)]*) and epithelia strongly promoted stromal proliferation (*K_eeo,esc_* >> *[EEOC(t)]* and *FX_eeo,esc_* >> 1) (Table 5). Therefore, molecules, like progestin, that decreased overall epithelial growth in monoculture (compared to experiments without that molecule) could still increase it in co-culture by inhibiting stromal cell proliferation.

Our model parameters suggest that the influence of each cell type on their own proliferation and on the proliferation of the other cell type differed between cells derived from different donors. For cultures generated from donor 271’s cells, our model predicted that epithelial cells strongly inhibited stromal cell proliferation and stromal cells slightly promoted epithelial cell proliferation. In contrast, donor 272’s epithelial cells strongly promoted stromal cell proliferation and stromal cells slightly inhibited epithelial cell proliferation.

The effects of the progesterone receptor inhibitor RU-486 and the cytokine IL-1*β* on overall epithelial and stromal cell density were also affected by epithelial-stromal regulation. The largest IL-1*β*-effect is seen for cells from donor 260 in co-culture (Fig. 7B,D), which also have the lowest value for *FX_il1b,eeo_* (0.327) (Table 5). Despite RU-486 strongly stimulating stromal cell death (high values of *FX_ru,esc_*) for donor 260’s and donor 272’s cells, the effects on the overall stromal cell density in mono- and co-culture were minimal because of the low stromal cell proliferation rates and low baseline stromal death rates (Fig. 6A, Table 5). Likewise, despite IL-1*β* strongly stimulating stromal cell death for cells from donor 272 (*FX_il1b,esc_* ∼ 10), its effect on overall stromal cell density is minimal in mono- and co-cultures (Fig. 7E-F).

### Differences in cell proliferation have minimal impact in IL-1β distribution in hydrogels that are less than 50 µL (post-swelling)

In 3D culture, mass transport through a hydrogel introduces the potential for diffusible molecules to form spatial gradients throughout the hydrogel (40,55–57). The diffusivity of the molecule, the rate of molecule uptake by cells, and cell population changes can all impact the distribution of exogenous molecules across a 3D culture system. Understanding how these parameters impact molecular gradients in a hydrogel is critical for the interpretation of experimental results from 3D culture.

We simulated the distribution of the largest (17 kDa) signaling molecule, IL-1*β*, added to the liquid culture media in the experiments used to calibrate our computational model (22). Smaller molecules would be expected to diffuse faster, thus having shallower gradients and less heterogeneity across the 3D culture. Endometrial epithelial and stromal cells both express IL-1*β* receptors (58,59), and these cell surface receptors mediate IL-1*β* uptake by cells (6). We used partial differential equations to create a reaction-diffusion model of IL-1*β* in the culture hydrogel. This model simulates the movement of IL-1*β* from the media into the cell-containing hydrogel matrix, including its diffusion and loss through internalization via receptors on epithelial and stromal cells (described in *the Methods*), and thus predicts the resulting molecular gradients and heterogeneity.

We first simulated the IL-1*β* distribution with the estimated parameter values (Table 4) for a constant cell density of 5⨯10^6^ epithelial cells/mL and 5⨯10^6^ stromal cells/mL within a 3 µL hydrogel, assuming a 6 µL steady state volume is reached after swelling prior to the start of the experiment. Note, a hemispherical hydrogel that is 6 µL post-swelling has a radius of 0.14 cm, which is the radius over which we solve our reaction-diffusion model. Our simulation predicted that IL-1*β* would saturate the co-culture gel (i.e. reach close to the media concentration of 1 ng/mL at all locations) within 5 hours and remain close to its saturated level (≥0.9 ng/mL) throughout the gel during the 20-day simulation. To check whether cell proliferation would affect the IL-1*β* distribution, we simulated a case with high epithelial and stromal proliferation in the presence of IL-1*β*: the “Estrogen, IL1*β*” experiments (mimicking an inflamed proliferative phase) conducted with cells from donor 272. Both the mono- and co-culture experiments begin with at least 5⨯10^6^ cells/mL and our ODE model predicted a ∼five-fold increase in total cell density in co-culture over the course of 15 days (Fig. 5B-C). Even with this high cell proliferation, our PDE model still predicted IL-1*β* to saturate the gel with a concentration of 0.9 ng/mL within 5 hours (Fig. 8A-C). This concentration is still greater than the apparent half maximal inhibitory concentration (IC50) for IL-1*β* inhibiting endometrial cell decidualization, 0.2 ng/ml (60), signifying that we may expect similar endometrial cellular responses at 0.9 and 1 ng/mL *in vitro*.

**Figure 8.**
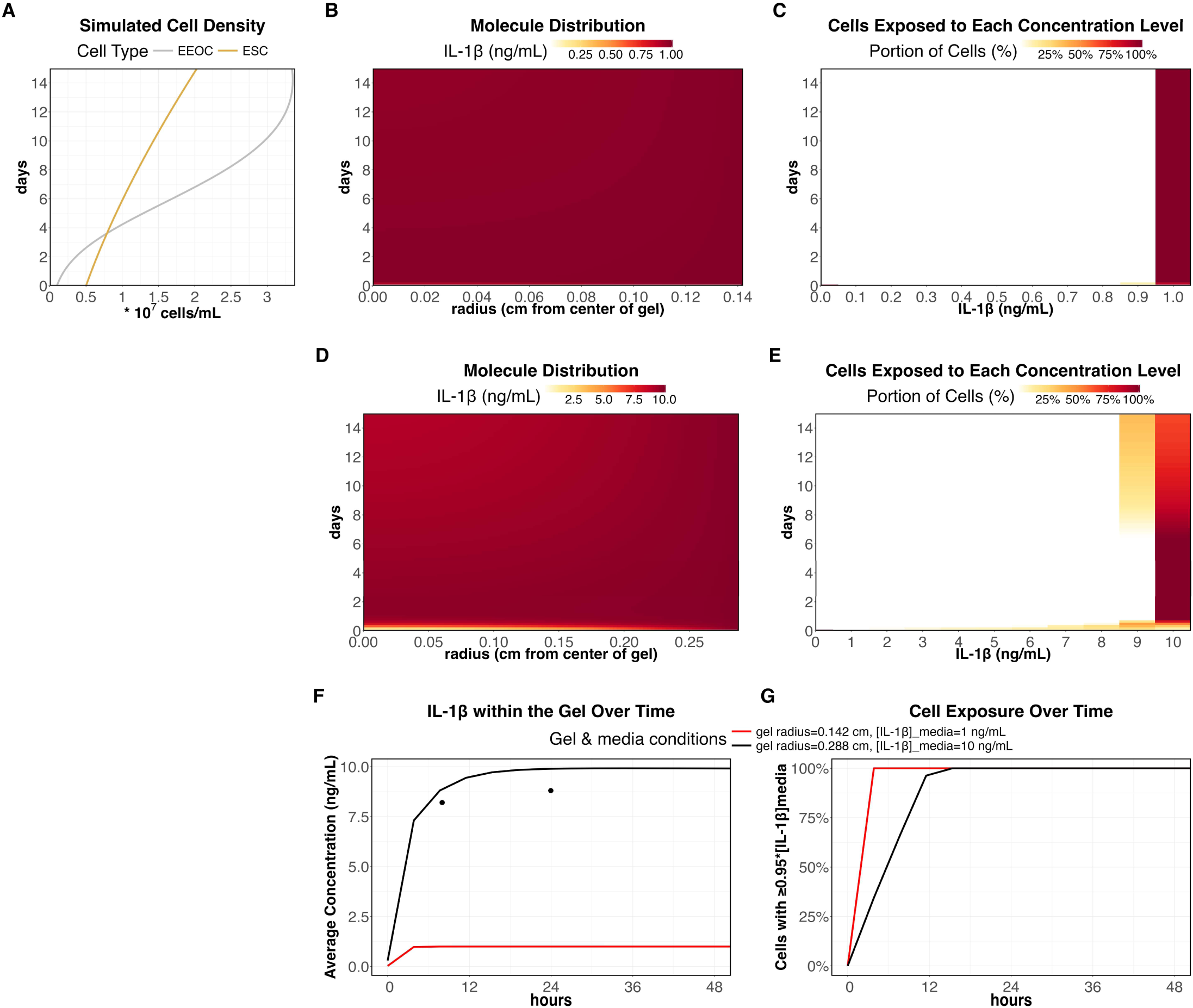
Simulated IL-1*β* gradients formed in 3D co-cultures. **(A)** Cell density predicted using our endometrial growth model (Eq. 2-3). **(B-E)** Simulated IL-1*β* distribution from the media into the gel using our PDE model with estimated parameter values in Table 4. We adjust these parameters to predict IL-1*β* distribution for co-cultures of similar size and added IL-1*β* as **(B-C)** Gnecco & Brown, et al. (2023) or **(D-E)** Valdez et al. (2017) (Table 6). For each simulation we show: (B, D) the IL-1*β* distribution across the gel, and (C, E) the portion of cells in the gel exposed to different levels of IL-1*β* (due to their location within the gel) is plotted as a histogram over time. The cell exposure histograms in C-E were calculated using 11 bins, ranging from 0 to the media concentration. In C-E, each x-axis label represents the middle concentration for each bin (e.g., “0.9” represents concentrations between 0.85 and 0.95 ng/mL). **(F)** The IL-1*β* concentration averaged across each gel at each timepoint. Here, we include experimental data for the larger gel used in Valdez et al. (2017). **(G)** The percentage of cells exposed to ≥0.95 times the media concentration of IL-1*β*.

**Table 6.** Comparison of methodologies in experiments used to generate this computational model of endometrial growth (Gnecco & Brown et al. (2023) (22)) and a previous study used in validation (Valdez et al. (2017) (21).

| Parameter | Value in Gnecco & Brown et al. (2023) | Value in Valdez et al. (2017) (IL-1β stimulation experiment) |
| --- | --- | --- |
| Hydrogel volume (pre-swelling) * | 3 μL | 25 μL |
| Media volume | 100 μL | 400 μL |
| IL-1β media concentration * | 1 ng/mL | 10 ng/mL |
| Media formulation | DMEM/F12<br>+ supplements for organoid expansion (e.g., rhEGF, ehNoggin, rhRspodin-1, etc)<br>+ estradiol (1 nM) | DMEM/F12/FBS |
| Gel base | 8-arm PEG vinyl sulfone (20 kDa) | 8-arm PEG vinyl sulfone (40 kDa) |
| Bioactive peptides included | Collagen I-derived peptide<br>Fibronectin-binding peptide<br>Basement membrane (collagen IV and laminin)-binding peptide | Fibronectin-mimetic peptide |
| Cell Types | Primary cells: endometrial stromal cells and epithelial glandular cells isolated from endometrial biopsies | Immortalized cells: endometrial epithelial cells (Ishikawa) and stromal cells (tHESC) |
| Cell-seeding density/method | 10<sup>7</sup> stromal cells/mL<br>10<sup>4</sup> epithelial organoids/mL | 4 × 10<sup>6</sup> stromal cells/mL<br>4 × 10<sup>6</sup> epithelial cells/mL |

\* = Parameter values that were set to specific values in different simulations to represent the equivalent experiment

This rapid IL-1*β* saturation in gels is consistent with results from a previous study which used a larger PEG-based hydrogel (25 µL pre-swelling, assuming 50 µL post-swelling) (21). We approximated this hydrogel as a hemisphere with a radius of 0.288 cm post-swelling, which is roughly twice the radius of the gel from the experiments used to calibrate our ODE model. In this larger hydrogel, endometrial epithelial cells and stromal cells were co-cultured, and the gel was separated from the media and later dissolved to recover intact cells and measure the concentration of molecules such as IL-1*β*. Following the addition of 10 ng/mL of IL-1*β* to the media, the concentration in this larger gel was 8 ng/mL within 9 hours and 9 ng/mL at 24 hours (21). We simulated the differences in gel size at steady-state and differences in media IL-1*β* concentration for these two studies (Table 6). Our simulation predictions for the larger gel were close to what was observed in this previous study (21); our model predicted that IL-1*β* reaches an average value of 8 ng/mL within 8 hours. Within 15 hours, IL-1*β* is fully saturated with ≥9 ng/mL throughout the gel (Fig. 8D-G).

Although we estimated model parameters using *in vitro* studies which included different cell types (see *Supplemental Methods*), we know that IL-1*β* diffusivity, receptor expression, and IL-1*β* cell uptake rate are gel- and cell-specific. However, limited data exists for these parameters for endometrial cells in 3D culture. We therefore reran our simulations considering ranges for these parameter values and using a small gel (6 µL, post-swelling) as a baseline (Table 4). As we varied each parameter, one-by-one, ten-fold from the baseline values in Table 4, we found decreasing diffusivity produced the largest gradients across the gel (Fig. 9A-B). Simulating the cell culture experiments from the study used to parameterize our ODE model (22), we saw similar predictions in gradient formation, signifying that the predicted difference in cell proliferation within these experiments is not enough to alter how much IL-1*β* is distributed for the assumed parameter values in Table 4 (Fig. 10A-B). Interestingly, our model predicted that differences in cell proliferation rates could lead to differences in IL-1*β* distribution for a larger hydrogel (Fig. 10C-D). These simulations suggest that smaller hydrogels may be more suitable for multi-day studies investigating signaling pathways within 3D cultures of proliferating endometrial cells.

**Figure 9.**
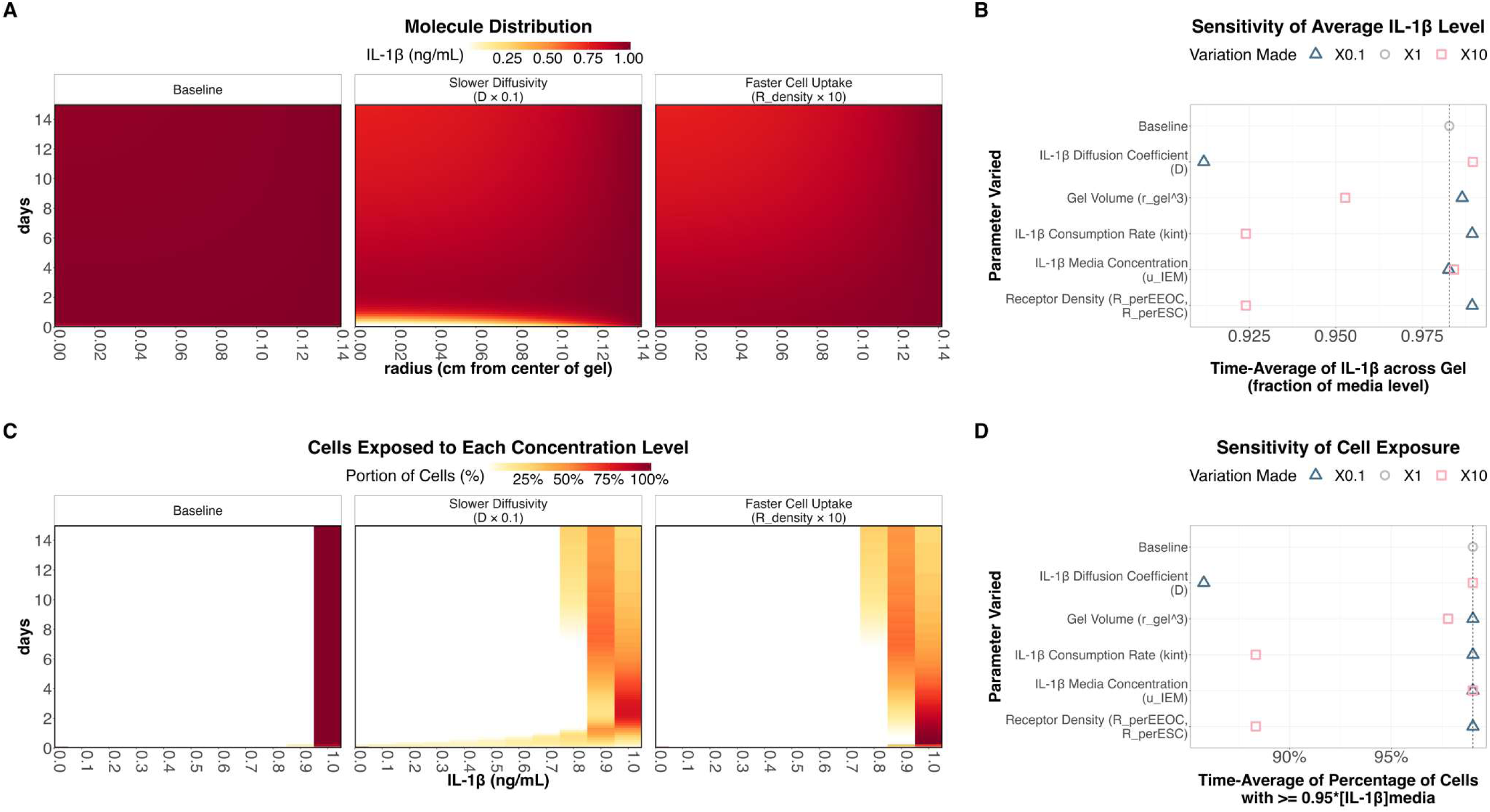
Sensitivity of predicted IL-1*β* distribution to gel- and cell- specific parameter values. Results from univariate sensitivity analysis, showing how each parameter variation affected the distribution of IL-1*β* throughout the hydrogel **(A, B)** and the IL-1*β* distribution transformed into a histogram of the percentage of cells at each concentration level **(C, D)**. For subplots A & C, cell density was simulated as that for donor 272, estrogen-IL1*β* co-culture case (Fig. 8A). In subplot C, R_density is used to denote a variation to the receptor density on EEOCs (R_perEEOC) and ESCs (R_perESC). For sensitivity analysis in B & D, each parameter was increased and decreased ten-fold from their baseline values in Table 4, and the result for no change in any parameter from baseline was plotted as a dashed line (also indicated by X1 result).

**Figure 10.**
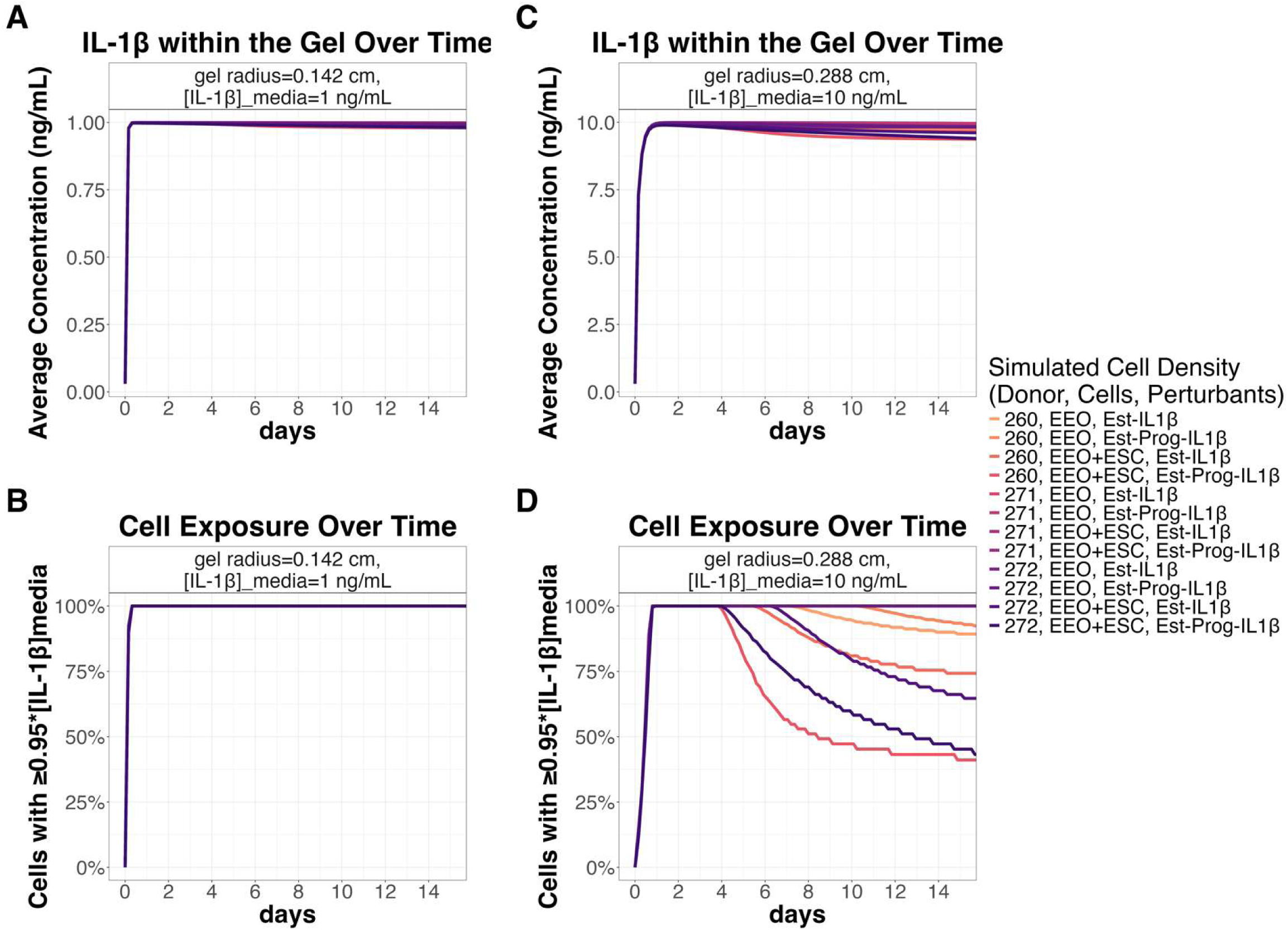
Sensitivity of predicted IL-1*β* distribution to cell growth. IL-1*β* reaction-diffusion was predicted for the time-course simulations of epithelial and stromal cell density in monoculture (EEO) and co-culture (EEO+ESC) experiments for different donors (260, 271, and 272) and perturbants (“Est”=Estrogen, “Prog”=Progestin, “IL1β”=Interleukin-1*β*), as shown in Figure 5. We show the resulting IL-1*β* distribution and cell exposure for a small gel (0.142 cm radius, 3 µL volume pre-swelling, 6 µL volume post-swelling) **(A-B);** and a larger gel (0.288 cm radius, 25 µL volume pre-swelling, 50 µL volume post-swelling) **(C-D).**

## Discussion

In this study, we created an ordinary differential equation (ODE)-based computational model of endometrial cell proliferation and calibrated this model using experimental results from a 3D *in vitro* model that was shown to mimic phases of the menstrual cycle. Because we modeled different cell types in mono- and co-culture, we were able to use this ODE model to compare how stromal and epithelial cells respond to hormones and cytokines and how cell-cell communication may contribute to observed differences in the growth of endometrial epithelial organoids in cultures generated using cells from different human donors.

Our model predicted that cell proliferation rates differed between cultures generated from different donors’ cells, as one would expect when working with primary cells. The model also predicted that the epithelial cell density in monocultures increased more overall than the stromal cell density in monocultures from the same donor. Looking next at the responses to progestin and IL-1*β*, there were some responses that were similar across donors and others that were donor-specific. In our model simulations, progestin consistently had little effect on stromal cell density in monocultures generated from each donors’ cells but had an even greater effect on stromal cells in co-culture. In contrast, both measurements and model simulations showed that progestin’s effect on EEO size varied in magnitude and direction depending on the donor (Fig. 7). Although the donor-to-donor differences observed in experimental measurements and our computational model simulations may stem from differences in disease phenotypes or other donor characteristics (Supplementary Table S1), there were not enough donors in this study to draw conclusions about the causes of these differences. It is still important to note, as a proof of principle, that the computational model can capture interindividual differences, which will facilitate the use of this computational model to understand a larger dataset.

The differences in both EEO and ESC responses to progestin in our mono- and co-culture simulations suggest epithelial-stromal crosstalk. Although few studies quantify changes in EEO size over time, one *in vitro* study noted that progestin neither inhibited nor stimulated the growth of EEOs generated from endometrial carcinomas and hypothesized that progestin responsiveness may be stroma-driven (61). Previous *in vivo* studies demonstrated stromal-epithelial crosstalk within the uterus of rodents, showing endometrial epithelial cells are necessary for the ESC decidualization response to progesterone (62) and that ESCs mediate progesterone-induced inhibition of endometrial epithelial cell proliferation (63).

In contrast to progestin, the endometrial cells from different donors responded more consistently to IL-1*β.* In our model simulations, IL-1*β* consistently increased the overall change in epithelial cell density and slightly decreased the overall change in stromal cell density across both mono-and co-culture experiments for all donors, compared to the experiments without IL-1*β* present. Given that the impact of IL-1*β* on epithelial cells was large and highly consistent in co-culture, this effect was likely through the action of stromal cells. Furthermore, our model predicted that IL-1*β* decreased the overall change in stromal cell density in both mono- and co-culture experiments, suggesting that stromal cells inhibit epithelial cell proliferation in the presence of estrogen and progestin; and that a reduction in this inhibition when IL-1*β* lowers stromal cell density led to a greater increase in epithelial density in co-cultures with IL-1*β* present (whereas epithelial cells in monoculture do not respond to IL-1*β* this way). In monoculture experiments, IL-1*β* decreased ESC proliferation and increased ESC apoptosis (22). In co-culture, IL-1*β* increased EEO size and epithelial proliferation, but changes to stroma were difficult to quantify in the presence of EEOs. Our computational model allows us to predict stromal cell density in co-culture, which aids our ability to infer how the presence of multiple cell types can affect endometrial tissue growth in response to different hormones and cytokines.

Both the progestin and IL-1*β* examples above demonstrate a role for cell-cell communication. Since the experimental study used to calibrate our ODE-based computational model is one of the first to create stable long-term co-cultures of primary endometrial epithelial and stromal cells by co-encapsulating them within a 3D hydrogel, it is difficult to compare such cell-cell effects in co-culture to previous studies. One study which co-encapsulated immortalized endometrial epithelial and stromal cells in a collagen gel found that epithelial cell monocultures had a lower DNA content than stromal cell monocultures without estradiol present; estradiol significantly increased the DNA content of epithelial cell monocultures (presumably through increasing cell density), decreased DNA content in stromal cell monocultures, and decreased DNA content in epithelial-stromal co-cultures (20). However, the use of immortalized cell lines in this experiment means the phenotype and interactions of these cells may differ from native tissues. Other studies used transwell plates or other means to culture stroma and epithelia in adjacent compartments. One such study (64) found that a larger percentage of primary ESCs were proliferating than endometrial epithelial cells. The addition of ESCs to endometrial epithelial cell cultures in another study increased the percentage of proliferating epithelial cells (5). However, these studies report percent proliferating cells and not changes in cell density, so they are not a direct reflection of the net effects of each cell type on the growth dynamics, as apoptosis may also be affected. Additionally, the use of a transwell co-culture model limits the study to probing paracrine signals between the two cell types, since physical interactions and mechanical signals cannot be transmitted through the transwell.

The 3D hydrogel matrix from the experiments used to calibrate our ODE model had the advantage of being able to co-encapsulate primary endometrial epithelial and stromal cells, allowing for physical interactions between the two cell types and mechanical signaling through cells interacting with the hydrogel matrix. However, mass transport in a 3D hydrogel can potentially allow for molecular gradients to form as molecules diffuse through the gel and interact with encapsulated cells. If significant gradients form within the hydrogel, then heterogeneity in cellular exposure to cytokines, growth factors, nutrients, and other molecules from the media may lead to heterogeneity in biological response. This should be considered while interpreting experimental readouts, particularly because heterogeneity of the responses of single cells cannot be characterized using results that are measured in bulk, such as analysis of media supernatants or bulk RNA sequencing.

To understand the potential for molecular gradients and heterogeneity in 3D culture, we developed a second, related model, using partial differential equations (PDEs) to model the distribution of a signaling molecule, IL-1*β*, through a hemispherical hydrogel containing proliferating cells. Our analyses showed that differences in cell proliferation rates yielded less spatial heterogeneity for IL-1*β* distribution in 6 μL (post-swelling) hydrogel cultures compared to larger (50 μL, post-swelling) cultures. This suggests that smaller cultures would be useful for comparing multiple cell types or cells from different donors because they provide more uniform distribution of both exogenous and endogenous bioactive factors like cytokines, regardless of variability in cell proliferation rates. Previous studies have shown that the presence of cells within the hydrogel could decrease oxygen and glucose at the hydrogel center over the course of one week (57). Another study found that over short time-scales (1 hour), cultures with more cells seeded had lower oxygen consumption rates, oxygen diffusion coefficients, and affinity towards oxygen compared to cultures with fewer cells seeded (65).

Unlike previous models, our PDE model predicts molecule distribution over multiple weeks, including mechanistic details like the expression of receptors on different cell types and their subsequent effect on molecule uptake by cells. By connecting our PDE model of molecular distribution to our ODE model of cell proliferation, we also consider how differences in the proliferative characteristics of endometrial cells (stemming from donors’ disease, menstrual phase, age, etc.) could impact the distribution and uptake of molecules in 3D environments. The potential for molecular gradients in 3D environments would also increase for larger molecules, e.g. antibodies used to investigate immune modulation as treatment for endometriosis. Inhibition of the IL-1/IL-33 signaling pathway has already been shown in a mouse model to produce smaller endometriosis lesions with fewer proliferating cells (66). There are ongoing clinical trials for a recombinant IL-1 receptor antagonist and a monoclonal antibody against IL-33 (67). Our computational modeling may benefit the development of assays and therapies for immune modulation. Indeed, ODE and PDE-based mechanistic models of molecule distribution have been used for other biomedical applications, such as in the design of cell implants for treating peripheral nerve disease (68) and in the design and analysis of hydrogels for directing blood vessel formation (69).

There are limitations to both our ODE-based and PDE-based computational models. Since we calibrated our ODE model of endometrial cell proliferation using data from only three donors, we have refrained from drawing conclusions regarding how the results here may stem from differences between the donors, e.g.: donor age; the menstrual phase tissues were taken at; and the disease diagnosis of each patient. We instead focused our discussion on identifying potential differences between cell types since there are many EEOs and ESCs in each experiment. The addition of more clinically-phenotyped donors may enable conclusions related to how differences in donor characteristics contribute to differences in endometrial cell proliferation. The present work serves as proof of principle that the modeling and parameterization can be done and can be used for future work with more donors.

Another potential limitation of our ODE model of cell proliferation is that it does simplify the growth of endometrial organoids by assuming constant epithelial cell height and spherical shape to make calculations. In the experimental study used to calibrate this computational model, researchers found that EEOC height was reduced in the presence of IL-1*β* and that EEOs sometimes became more ovoid (22). Using H&E staining and 3D image analysis of Epcam-vimentin staining, they were able to deduce that cell height would change but the number of epithelial cells within an organoid was unaltered, suggesting that modeling changes to cell height here was unnecessary, but this may not hold under all conditions. We assume spherical shape using the longest radius for each EEO, which may slightly overestimate the number of cells in ellipsoidal EEOs. Automation of 2D and 3D image analysis of 3D cultures may make quantitative studies faster, more robust, and capable of accounting for perturbant effects on morphology.

One additional consideration is that on average ∼11 EEOs were measured in each experiment, and we used the average EEO size trajectory to calibrate our ODE model. The ODE model parameters are thus capturing the average effect of estrogen, progestin, RU-486, and IL-1*β* on EEO size and not variability within an individual donor’s cells. In addition, we model the effects of these molecules and cell types as constant effect terms that depend on their presence or absence and not the exact concentration available nor length of exposure. Although this was a suitable assumption for molecules maintained at a constant level within the media (e.g., estrogen, progestin, and IL-1*β*), we could explore density-dependent functions for the pro-proliferative effects of cell types present in the future.

Although we were able to optimize parameters for both epithelial and stromal cells, our optimization included more data for epithelial cells than stromal cells. As described in the *Methods*, our optimization included measurements of EEO radius at least 5 timepoints per experiment conducted with each donor’s cells in mono- and co-culture. In contrast, our optimization only included one end-point measurement from Live/Dead staining per experiment for monocultures generated from a single donor. Since the optimization identifies parameters that minimize the error between our model simulations and the experiment measurements, having this tenfold difference in data-density effectively means that minimizing the errors associated with EEOs becomes ten times more important to the algorithm than minimizing the errors associated with ESCs. Therefore, our optimization results can be interpreted as primarily identifying the ESC proliferation and death rates that allow for our simulations of EEO size in mono- and co-culture to match the time-course measurements.

Lastly, the simplifying assumptions made for our PDE model of IL-1*β* distribution are discussed in the *Methods* section and include assuming angular symmetry and equilibrium swelling conditions, assumptions appropriate for this static 3D culture system that was sustained for two-week experiments. Although we assume a constant value for the diffusion coefficient during experiments, fibroblasts have been shown to deposit collagen which can lead to hydrogel contraction and thereby change the diffusivity through the gel (52,70). The effects of cells on gel structure and diffusivity as they are cultured for multiple weeks should be explored more in future experimental and computational studies.

In conclusion, we provide here two computational models; one an ODE-based model of cell proliferation in 3D culture, and the second a PDE-based diffusion-reaction model of a bioactive signal added to the media of 3D cultures. These models can be re-calibrated using data from different experiments and research groups to study multicellular systems, to optimize cell culture conditions, and ultimately to further our ability to research and treat gynecological pathologies.

## Supplementary Materials

### Supplementary Materials and Methods

#### Estimating Parameters for IL-1*β* Model from the Literature

To summarize how these parameters were identified: the hydrogel radius and IL-1*β* media concentration are defined by the experimental conditions used by Gnecco & Brown *et al.* (2023) (22). We used the average of reported K_D_ for IL-1*β* to IL-1R1 and IL-1R2 (0.52 nM and 3.8 nM, respectively (51)). The diffusion coefficient for IL-1*β* through the cell-containing PEG hydrogel in (22) has yet to be measured, so we explored a range here. As a benchmark we expected molecules moving through a cell-containing gel to be 30% slower than their diffusivity in media. This was estimated by Kihara et al. (2013) using molecules ranging from 3 – 40 kDa moving through a soft collagen gel (∼0.2wt%) containing ∼10^5^ stromal cells/mL (52). Using the Stokes-Einstein equation, we estimated the diffusion coefficient for IL-1*β* in media to be 7.4⨯10^-7^ cm^2^/s. Since Gnecco & Brown *et al.* (2023) used a stiffer hydrogel (3wt%) that contains more cells (∼10^7^–10^9^ cells/mL), we predicted the diffusion coefficient of IL-1*β* through their gel to be 5⨯10^-7^ cm^2^/s, but we explored a range from (5⨯10^-8^ cm^2^/s – 5⨯10^-6^ cm^2^/s). Receptor expression and receptor internalization rates are highly cell-type dependent and may change based on a cell’s microenvironment. Tabibzadeh et al. (1990) performed a Scatchard analysis on human endometrial epithelial glands in 2D culture and found they expressed ∼4.55 fmol of IL-1 receptors/mg of protein (54). To estimate the number of IL-1 receptors per cm^2^ of the cell surface area we assumed that: each cell contains 7⨯10^-7^ mg of protein (71); only 50% of an epithelial cell’s surface was available for binding in a previous study since it was conducted in 2D (54). Assuming an epithelial cell radius of ∼6 µm, we estimated the density of IL-1 receptors to be 8.5⨯10^8^ receptors/cm^2^. To calculate the number of receptors per EEOC, we needed to first estimate the percent of epithelial surface area available for binding in 3D culture. By dividing the EEO surface area by the density of cells, we estimated ∼20% of an epithelial cell’s surface would be available for binding when they aggregate and form organoid structures in 3D. This receptor expression level and surface area availability yields 800 IL-1 receptors per EEOC and 1700 IL-1 receptors per ESC (assuming stromal cell radius of 4 µm with 100% of a stromal cell’s surface available for binding). We explored a range of receptor expression from 8.5⨯10^7^ – 8.5⨯10^9^ receptors/cm^2^. Lastly, the receptor internalization rate ranges from 2 – 6 ⨯ 10^-4^ (53,72), so we explored the wider range of 10^-5^ – 10^-3^ s^-1^ in this analysis.

### Supplementary Tables and Figures

**Table S1.** Characteristics of the donors of the endometrial biopsy samples that were used to generate experimental 3D culture models.

| **Donor ID** | **Age** | **Menstrual Cycle Phase** | **Diagnoses** | **Has Regular Menstrual Cycles** |
| --- | --- | --- | --- | --- |
| 260 | 27 | Late secretory | Endometriosis | yes |
| 271 | 36 | Secretory | Fibroids | yes |
| 272 | 42 | Proliferative | Abnormal uterine bleeding | no |

**Figure S1.**
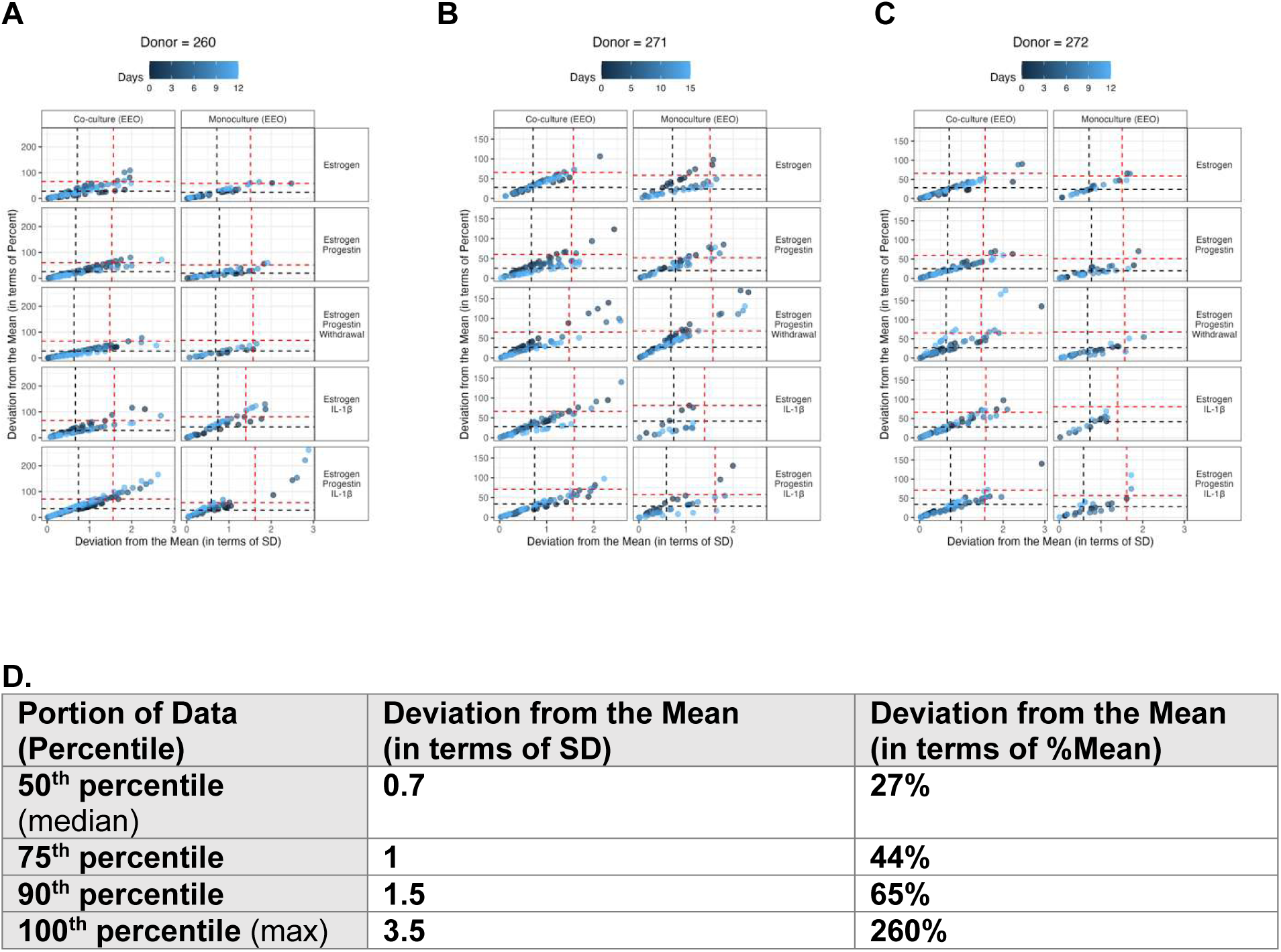
Experimental data used to set epithelial growth gate following optimization. We calculated the deviation from the mean of organoid measurements at each time-point for each experiment and donor. **(A-C)** The deviation of each organoid measurement from the mean in terms of standard deviation (SD) and percent deviation (%Mean) for each experiment and donor source at each time point. The black and red dashes in **A-C** represent the 50^th^ percentiles and 90^th^ percentiles, respectively. **(D)** Summary table across all donors.

**Figure S2.**
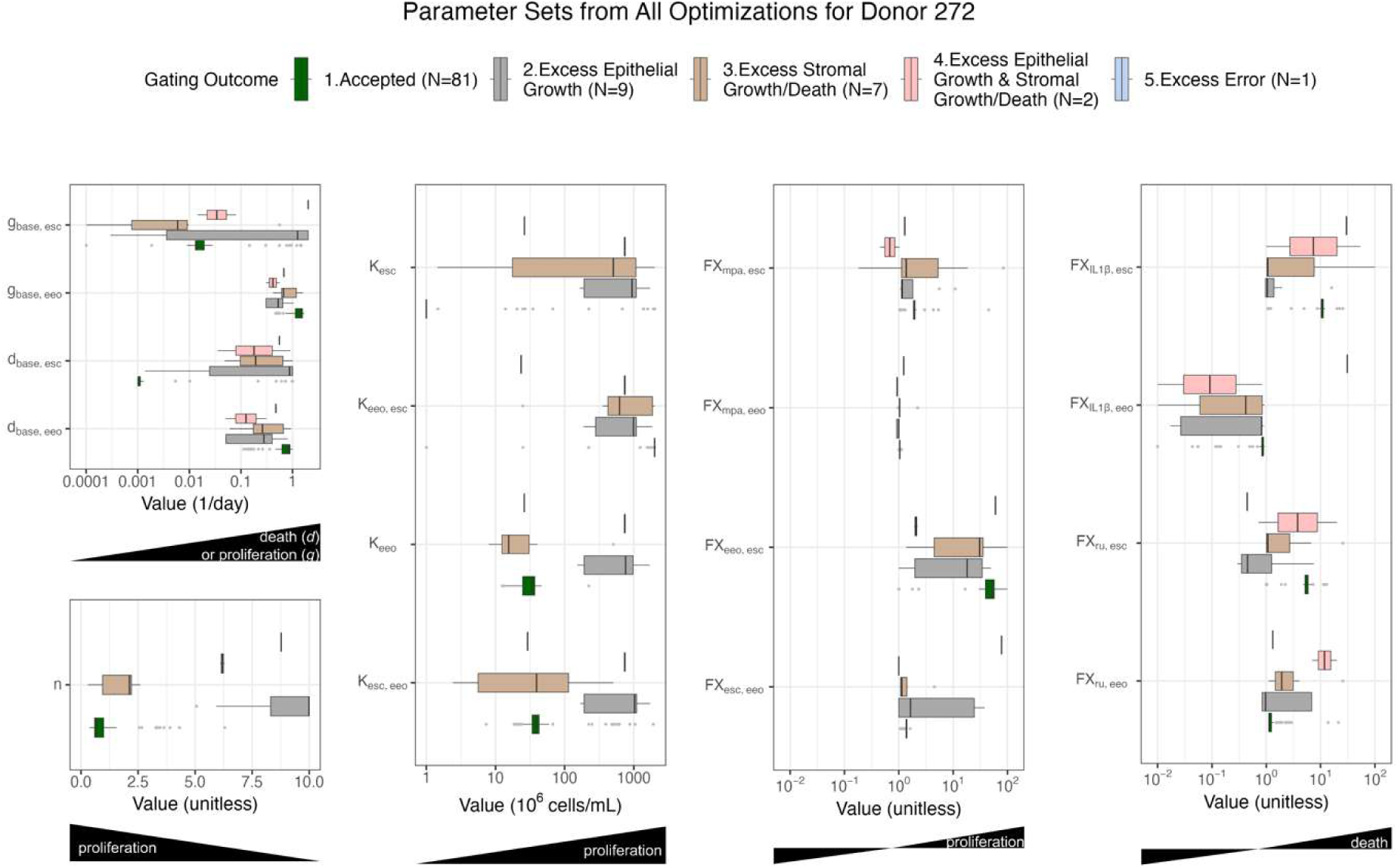
Gating optimization solutions by performance and biological relevance for donor 272. The distribution of optimized parameter sets. The corresponding simulations are shown in Figure 2.

**Figure S3.**
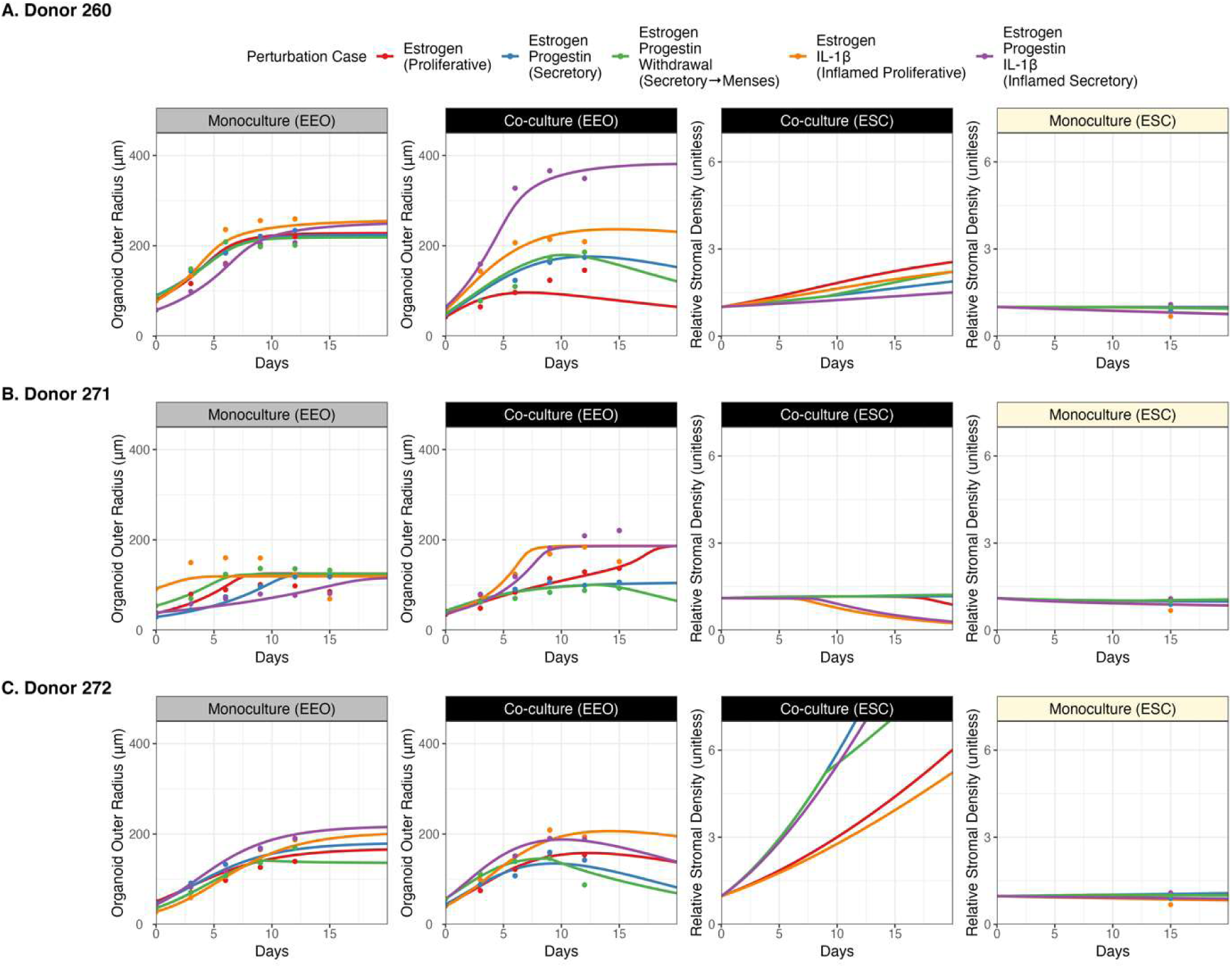
Comparison of simulations using the best accepted optimization of model parameters for each donor. The model was optimized using data from experiments conducted with cells from donors, including: **(A)** donor 260, **(B)** donor 271, **(C)** donor 272. For each donor, we show simulations for each of the fifteen experimental conditions (five perturbations ⨯ three cell culture scenarios) using the parameter set with the least sum-squared optimization error. The cell culture cases for experiments include: EEO monoculture (left-most plot), the co-culture of EEOs with ESCs (data for each of the center two plots), and the ESC monoculture (right most plot). The “best” accepted optimization is the parameter set that produced simulations with the lowest sum-squared error (Eq. 7) out of the parameter sets that were accepted according to the criteria described in *Methods*.

**Figure S4.**
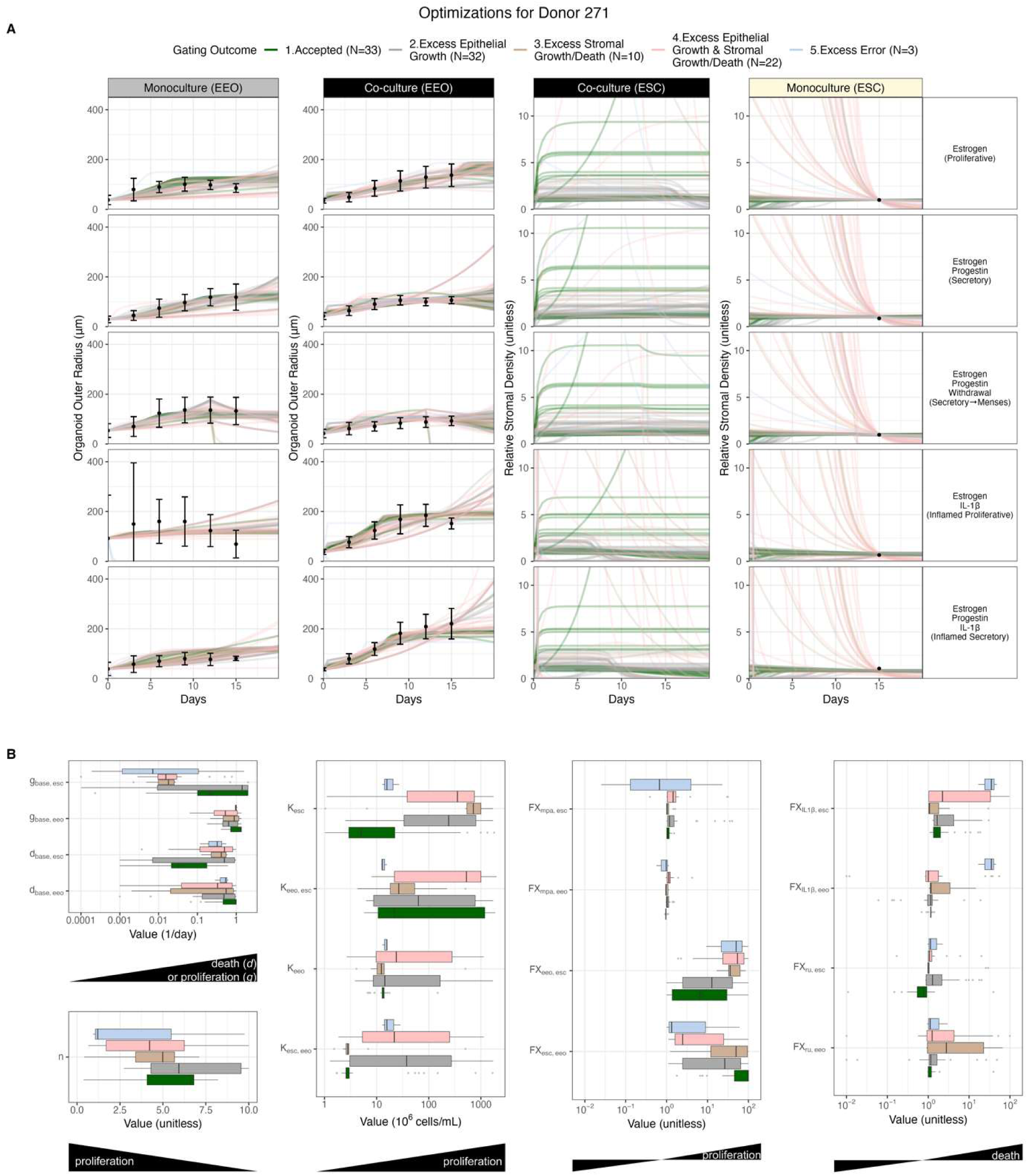
Gating optimization solutions by performance and biological relevance for donor 271. **(A)** Simulations (lines) and experimental data (dots), for all 15 experimental conditions as described by the column and row titles, similar to the format presented in Figure 2. **(B)** Corresponding distributions of optimized parameter value sets.

**Figure S5.**
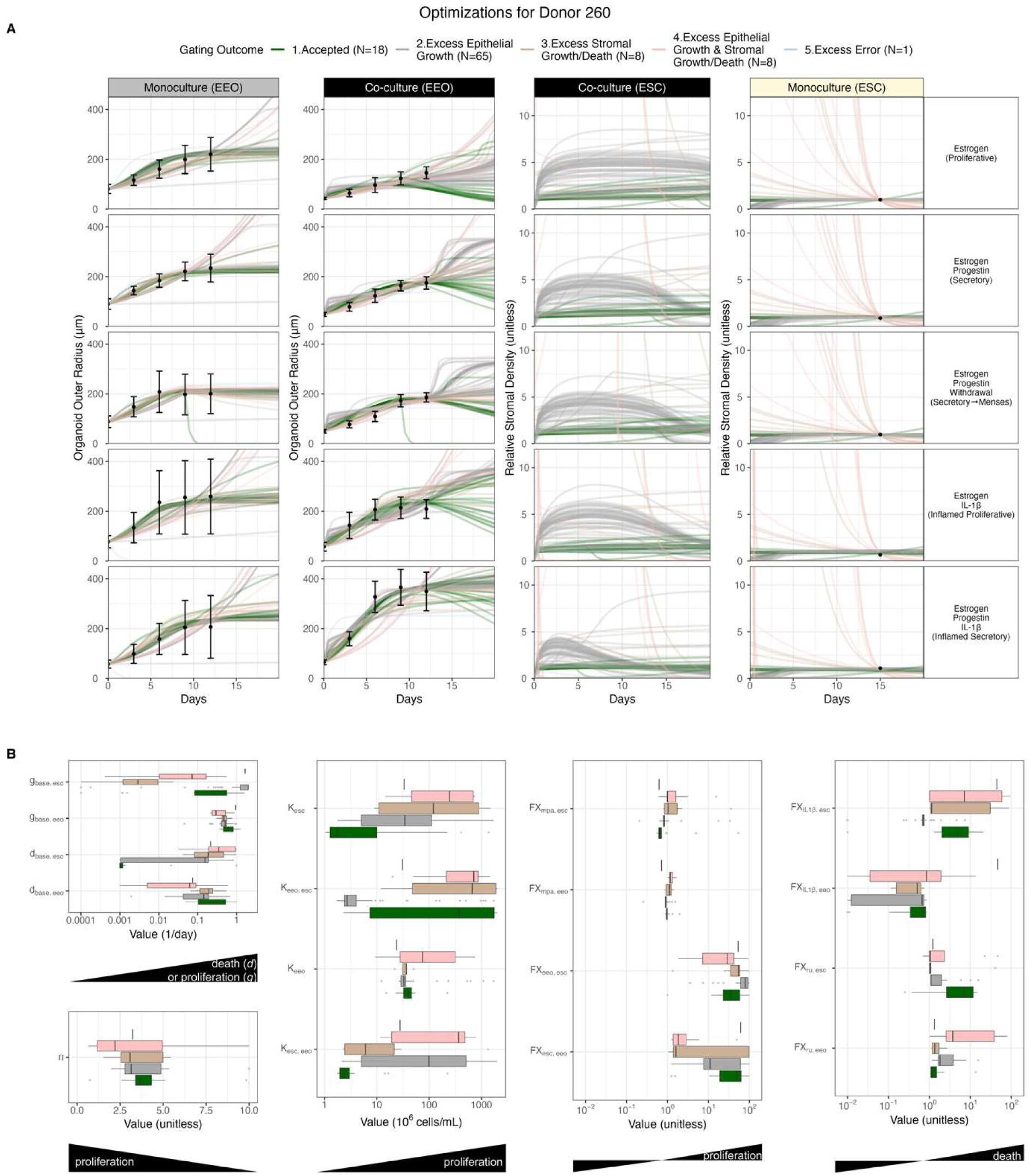
Gating optimization solutions by performance and biological relevance for donor 260. **(A)** Simulations (lines) and experimental data (dots), for all 15 experimental conditions as described by the column and row titles, similar to the format presented in Figure 2. **(B)** Corresponding distributions of optimized parameter value sets.

**Figure S6.**
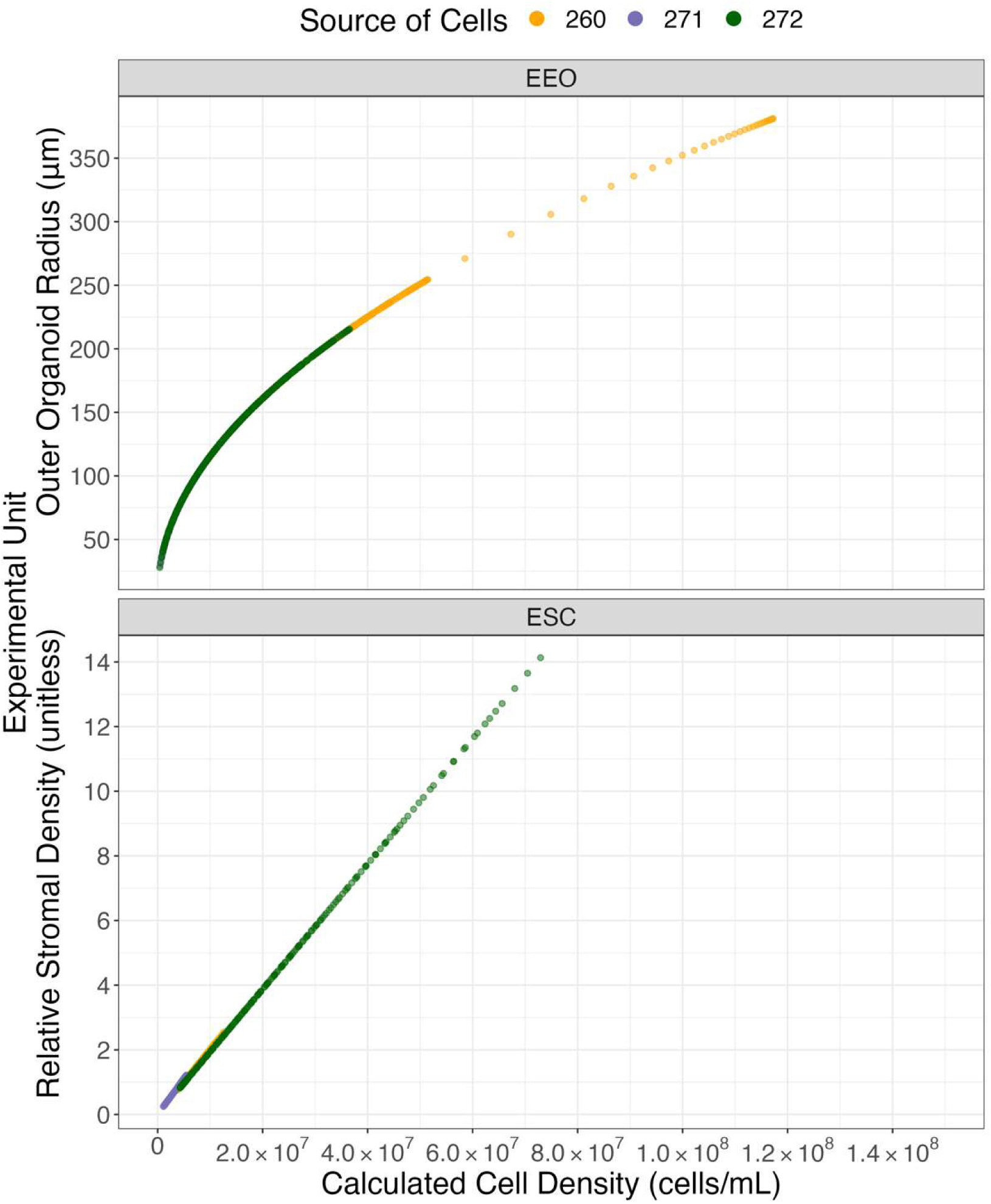
Converting between experimental units and cell density for simulations. **(A)** Simulated EEO outer radius (µm) vs. its converted cell density value for each donor, calculated using Eq. 4. **(B)** Simulated relative ESC density vs. its converted absolute cell density, calculated using Eq. 6. Experimental data was collected from experiments that used cells from different donors.

